# Infraslow histaminergic dynamics govern priming states to gate moment-to-moment memory accessibility

**DOI:** 10.1101/2025.11.13.687922

**Authors:** Yoshikazu Morishita, Yuki Takamura, Kyoka Nishimura, Yuto Yokoi, Yuya Ishihama, Rentaro Idutsu, Misato Ono, Reika Matsumoto, Natsuko Hitora-Imamura, Masabumi Minami, Hiroshi Nomura

## Abstract

Memory expression fluctuates even in response to identical cues, suggesting that ongoing brain states bias memory accessibility. However, the cellular and circuit principles governing these state-dependent fluctuations remain unclear. Here, we show that spontaneous pre-cue activity of histaminergic neurons in the hypothalamic tuberomammillary nucleus (TMN) modulates the expression of reward-associative memory in mice. TMN histaminergic activity exhibited infraslow dynamics (0.05–0.1 Hz) that closely tracked an integrated brain–body state. Closed-loop cue delivery during high histaminergic states enhanced memory expression. Brief optogenetic activation or inhibition of these neurons before the cue bidirectionally modulated memory expression, and direct activation of histaminergic terminals in the basolateral amygdala (BLA) was sufficient to enhance memory expression. Furthermore, histaminergic inhibition before the cue impaired the cue-evoked BLA population response. Thus, ongoing histaminergic activity exerts an infraslow, state-setting influence that primes BLA circuits for robust cue responses, and in turn, modulates moment-to-moment memory accessibility.

## INTRODUCTION

Memory accessibility fluctuates on the scale of seconds to minutes even during sustained wakefulness, as in the familiar “tip-of-the-tongue” state^1^ and sudden, spontaneous retrieval of a memory^2^. This observation suggests that transient internal brain states gate information flow and shape cognitive output in real time^3–5^. Clinically, aberrant state transitions are evident as fluctuating cognition in Lewy body dementia^6^, peri-ictal memory impairment in temporal lobe epilepsy^7^, and intrusive memories in post-traumatic stress disorder^8^. Although global physiological and psychological states (e.g., arousal, mood, metabolic status) are known to influence cognitive performance^9–12^, the cell types and circuit principles that generate moment-to-moment state transitions remain undefined. This gap impedes mechanistic understanding of maladaptive state dynamics observed across neurological and psychiatric disorders^13^.

Any candidate system should (i) have access to core memory circuits and be positioned to regulate the local network dynamics of those circuits, (ii) be causally involved in cognitive function, and (iii) operate on seconds-to-minutes timescales during wakefulness. The histaminergic system, originating in the tuberomammillary nucleus (TMN), broadly innervates key memory structures, including the cerebral cortex, hippocampus, and amygdala^14,15^, where histamine receptors regulate neuronal excitability and synaptic transmission^16,17^. Furthermore, perturbations of this system have been linked to memory impairments, from the neuronal loss in neurodegenerative conditions like Alzheimer’s disease to the side effects of common antihistamines^16,18–21^. Conversely, enhancing histaminergic transmission has been shown to promote the expression of otherwise inaccessible memories^22,23^. However, prior work has largely framed histaminergic activity within the binary sleep–wake cycle^24,25^ and has relied on exogenous manipulations that lack sufficient temporal precision. This focus does not readily account for within-wakefulness fluctuations in memory accessibility on seconds-to-minutes timescales. This raises the possibility that a previously unrecognized, more dynamic mode of histaminergic signaling during wakefulness contributes to these fluctuations.

Here, using cell-type–specific recordings and temporally precise manipulations in behaving mice, we test whether a dynamic within-wakefulness mode of histaminergic neuronal activity biases moment-to-moment memory accessibility. We first reveal that this activity is characterized by infraslow (0.05–0.1 Hz) fluctuations that closely track an integrated brain–body state (e.g., EEG, facial movements, and pupil dynamics). We then demonstrate that these fluctuations causally gate memory accessibility through gain-like modulation of cue-evoked population responses in the basolateral amygdala (BLA). These findings establish the principle that infraslow neuromodulatory dynamics generate internal brain states that gate cognitive function, providing a circuit-level basis for moment-to-moment memory accessibility in health and disease.

## RESULTS

### Infraslow histaminergic dynamics closely track an integrated brain**–**body state

We monitored the activity of TMN histaminergic neurons in HDC-IRES-Cre mice expressing GCaMP6s using fiber photometry (Figures 1A, 1B, and S1). In head-fixed mice without external stimuli, these neurons showed spontaneous oscillations with a clear peak in the 0.05–0.1 Hz range and exhibited strong bilateral TMN synchrony (Figures 1C, 1D, and S2A–S2C). We next examined how this activity related to cortical EEG activity and facial dynamics. Cross-correlation analysis revealed that the gamma-band power (30–80 Hz) and several facial movements consistently preceded histaminergic neuronal activity by 0–1 s, whereas very-low-frequency (VLF) EEG (0.01–0.5 Hz) peaked several seconds later (Figures 1E and 1F). To further test whether these signals predict histaminergic dynamics, we constructed a cross-validated linear encoding model that used the 2-s recent history of EEG signals, pupil dynamics, and facial features (Figure S2F). The full multimodal model achieved higher cross-validated R^2^ than any single-feature model and outperformed circularly shifted controls (Figures 1G and 1H). Block-shift permutation (ΔR²) identified nose movements and gamma-band power as principal predictors (Figure S2G). The predicted activity also reproduced a prominent 0.05–0.1 Hz peak, which was not evident in most individual signals (Figures 1D, 1I, S2D, and S2E). Together, these findings indicate that histaminergic neurons exhibit infraslow fluctuations tightly coupled to an integrated brain–body state, defined as a composite of EEG bands, pupil, and orofacial features.

**Figure 1.**
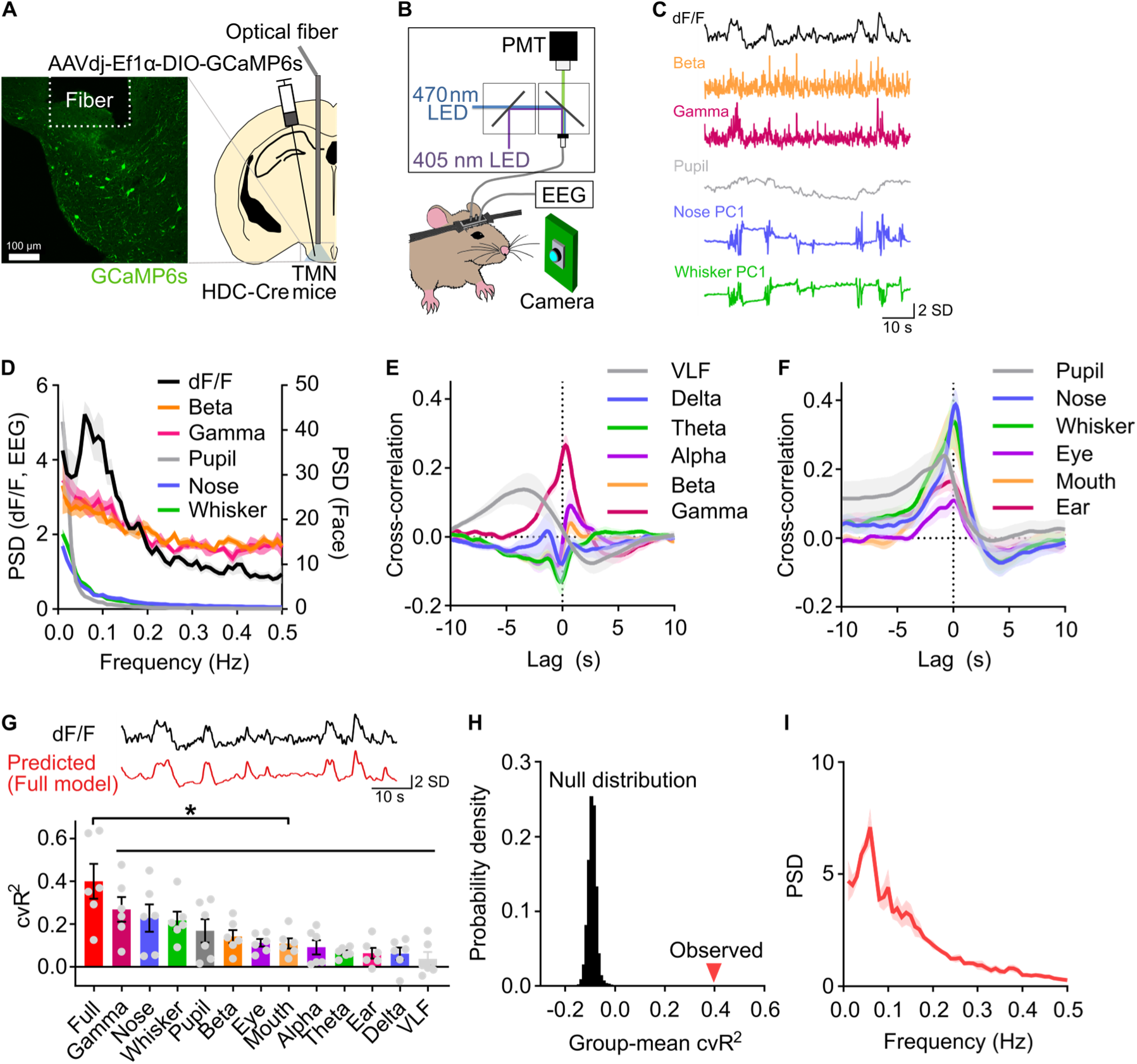
Infraslow histaminergic dynamics closely track an integrated brain–body state. (A) Schematic of viral injection and fiber placement for fiber photometry in histaminergic neurons, with a representative image showing GCaMP6s expression in the TMN. (B) Schematic of the simultaneous recording setup. (C) Representative traces showing spontaneous histaminergic activity, EEG band-limited amplitude envelopes, and facial-movement dynamics in a head-fixed mouse. (D) Normalized power spectral density (n = 6 mice). (E and F) Cross-correlation function between histaminergic neuronal activity and EEG-band power (E) and between histaminergic activity and facial motion energy (F). Positive lags indicate that EEG/facial signals lead the histaminergic activity. (G) Top: Example trace of recorded histaminergic activity and its prediction by the linear encoding model. Bottom: The cross-validated coefficient of determination (cvR^2^) for the full model and models using only single feature groups (*P < 0.05, one-tailed paired t-test with Holm-Bonferroni correction). (H) The observed group-mean cvR^2^ of the full model compared against a null distribution generated by shuffling the data (P = 1.0 × 10^-5^, Permutation test). (I) PSD of the predicted histaminergic activity from the full model. Data are mean ± SEM.

### Pre-CS histaminergic neuronal activity is enhanced in trials with high levels of memory expression

To test the relationship between histaminergic neuronal activity and memory expression, we used a trace appetitive Pavlovian conditioning task, in which anticipatory licking provides a reliable, trial-by-trial index of memory expression^26,27^. In this task, mice underwent conditioning in which a sucrose solution (unconditioned stimulus, US) was presented 1 s after a 2-s tone (conditioned stimulus, CS) (Figure 2A). After several days of conditioning, the mice exhibited predictive licking responses to the CS (Figure 2B), indicating successful expression of tone-reward memory. However, licking rates varied among trials, even after the mice learned about the association (Figure 2C). We defined high– and non-expression trials based on the licking rate during CS–trace period (the 3-s window from CS onset to US onset) (Figures 2D and S3A). Trials with licking rates in the top 10% were labeled as high-expression trials. To control for potential satiation effects and focus on memory expression failure, the first 10% of non-licking trials were chosen as non-expression trials. Baseline motivation was comparable between trial types, as indicated by similar pre-CS licking rates (Figure 2D). The median number of trials and intervals between trials did not differ between the two types of trials (Figures S3B and S3C). We then compared the activity of histaminergic neurons in the high– and non-expression trials and found that the activity was higher during the CS–trace period in the high-expression trials than in the non-expression trials (Figures 2E and 2G). Furthermore, the activity was already elevated before CS onset (Figures 2E and 2F). Given that the intervals between trials varied randomly between 23 and 29 s, the CS onset was unpredictable. Variation in licking rates between the high– and non-expression trials was not related to differences in arousal, because pupil size and its derivative before CS onset did not differ between the two trial types (Figures 2H and S3I). These results suggest that increased histaminergic neuronal activity preceding CS onset is associated with memory expression.

**Figure 2.**
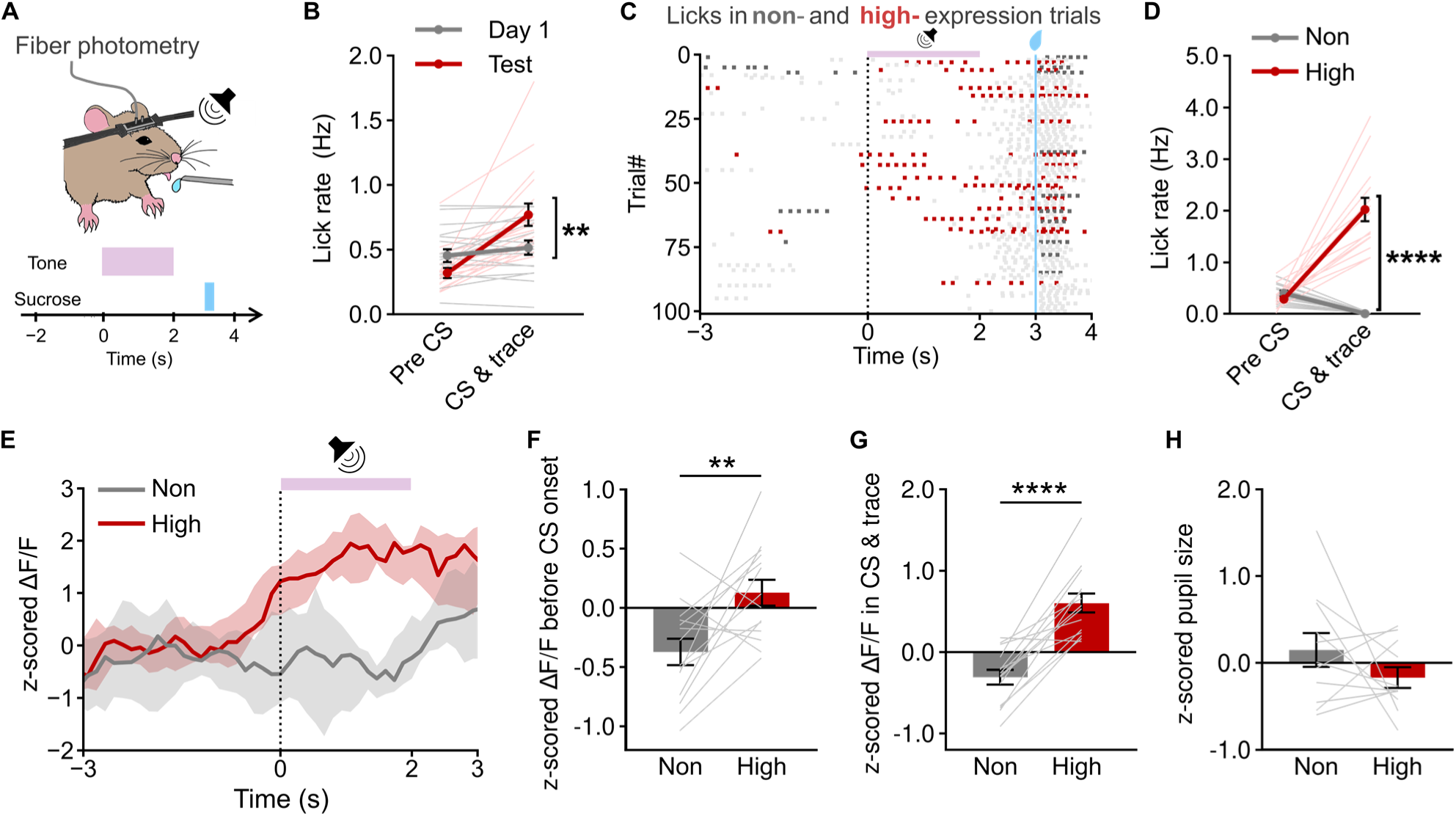
Histaminergic neuronal activity is elevated before CS onset in trials with high levels of memory expression. (A) Tone-reward conditioning procedure: mice received a sucrose solution (unconditioned stimulus, US) 1 s after a 2-s tone (conditioned stimulus, CS). (B) Licking rate during 3 s pre-CS period and CS–trace period (Day 1 vs. Test, n = 14 mice). (C) Raster plot of licking from a representative mouse (red, high-expression trials; gray, non-expression trials; light gray, other trials). (D) Licking rates during pre-CS and CS–trace periods. (E) Representative histaminergic activity (median ± top and bottom 25%). (F) Pre-CS activity across all mice. (G) Mean activity during the CS–trace period across all mice. (H) Pupil size before CS onset (n = 11 mice). Data are mean ± SEM unless noted.

It could be argued that pre-CS histaminergic neuronal activity reflects licking behavior itself or the motivation to lick, rather than memory expression; however, our results do not support this possibility. To test this, we analyzed activity on the first conditioning day. The licking rates were similar between the CS–trace and pre-CS periods (Figure 2B), suggesting that the mice had not yet formed a tone-reward association. Despite this, licking rates varied between trials, possibly reflecting different motivations or other internal states (Figure S3D). We compared trials with more licks and those with no licks (Figures S3D and S3E) and found that histaminergic neuronal activity before CS onset was similar in both trial types (Figure S3G). In contrast, activity during the CS–trace period was higher in trials with more licks than in trials with no licks (Figure S3H). Thus, pre-CS histaminergic neuronal activity is specifically associated with successful memory expression, while activity during the CS–trace period is partly related to licking behavior.

### CS presentations during high histaminergic activity enhance memory expression

To directly test whether high histaminergic states facilitate memory expression, we implemented a closed-loop system that delivered the CS–US pair when the photometry signal met predefined high– or low-state thresholds (Figure 3A). Trials were pre-randomized to high or low states. CS–US was triggered only when ongoing activity matched the assignment, ensuring balanced trial counts and inter-trial intervals (Figures S4A and S4B). Post hoc analysis verified that CS onset typically occurred within 2 s of threshold crossing (Figures 3B and 3C). In mice trained on the tone-reward association, licking during the CS–trace period was higher in high-state than in low-state trials (+39.0 ± 9.0%) (Figures 3D and 3E). In contrast, pre-CS licking rates were similar between conditions (Figure S4C). In the absence of CS, licking did not differ between the high– and low-state epochs (Figures S4D–S4F). Thus, a heightened histaminergic state enhances memory expression.

**Figure 3.**
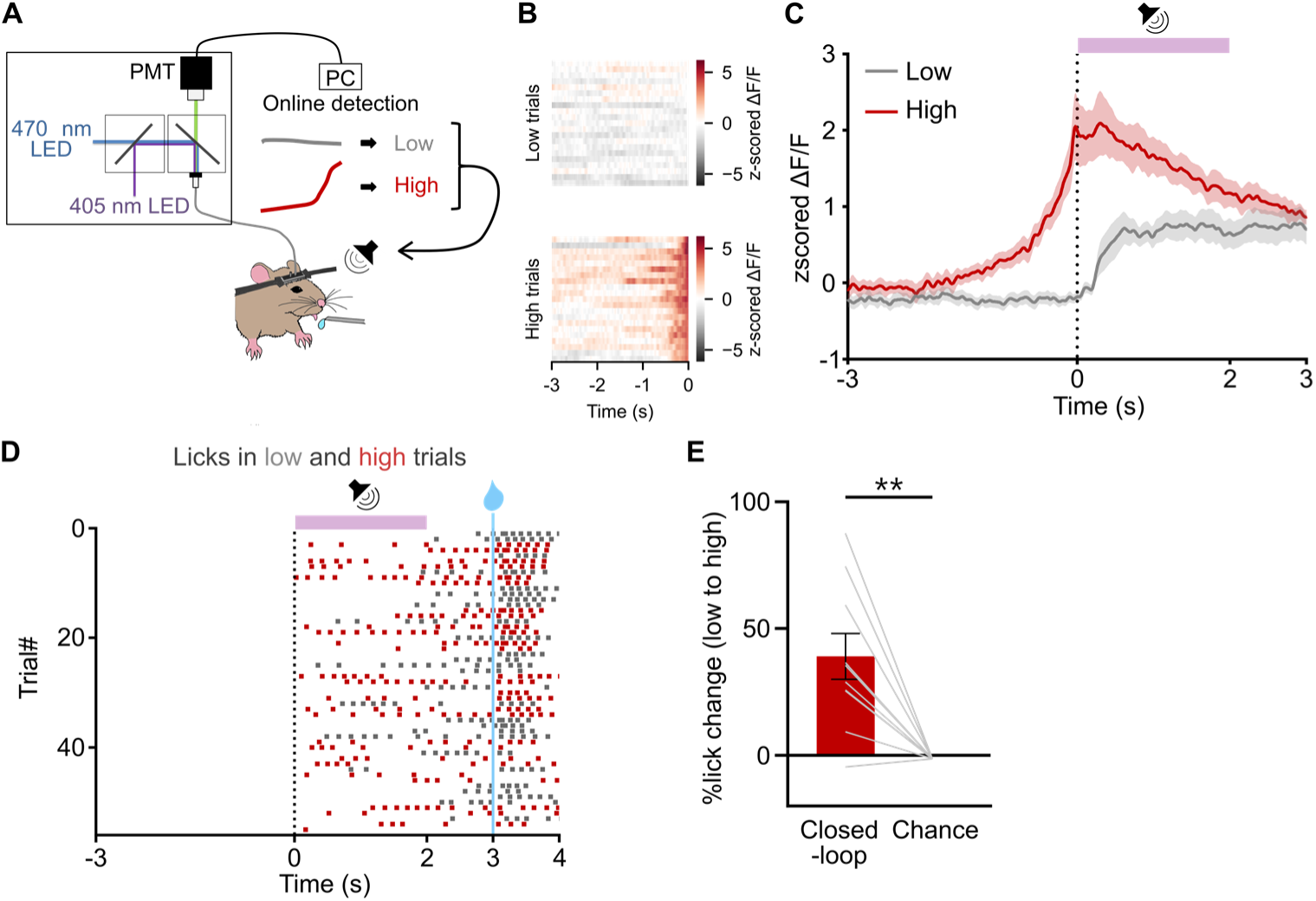
Memory expression is enhanced when a closed-loop system provides CS based on an increase in histaminergic neuronal activity. (A) Schematic of the closed-loop system for CS–US presentation triggered by the activity of histaminergic neurons. The CS–US was presented either when the activity exceeded an upper threshold (high) or when it remained below a lower threshold for 10 s (low). (B) Heatmap showing histaminergic neuronal activity in a representative mouse during high and low trials. (C), Histaminergic neuronal activity around the CS presentation in high and low trials (n = 10 mice). (D), Raster plot of licking events in high (red) and low trials (gray) for a representative mouse. (E), Percentage increase in the lick rate in high trials compared with low trials (**P = 0.0039, Wilcoxon signed-rank test, n = 10 mice). Data are mean ± SEM.

### Pre-CS inhibition of histaminergic neuronal activity reduces memory expression

To test for a causal link between histaminergic neuronal activity and memory expression, we optogenetically inhibited these neurons. We expressed Cre-dependent halorhodopsin (eNpHR) in TMN histaminergic neurons of HDC-IRES-Cre mice and implanted bilateral optical fibers (Figure 4A and S5). After trace appetitive Pavlovian conditioning, we delivered a green light to the TMN for 4 s (−1 to +3 s from CS onset). Memory expression was quantified using the area under the receiver operating characteristic curve (auROC) comparing pre-CS and CS–trace licking on each trial^28^. We found that light delivery reduced auROC in the eNpHR group, but not in the EYFP control group (Figures 4B and 4C). Brief inhibition restricted to −1–0 s similarly reduced the auROC (Figures 4D and 4E), whereas post-CS (0–3 s) had no effect (Figures 4F and 4G). Although predictive licking rates could be influenced by changes in motivation, auditory processing, attention, or arousal, these possibilities were ruled out because the optogenetic inhibition did not affect spontaneous licking (before sucrose presentation; Figure 4H), sucrose-induced licking (Figure 4I), acoustic startle response (Figure 4J), prepulse inhibition (Figure 4K), or pupil size (Figure 4L). Thus, pre-CS histaminergic neuronal activity supports the expression of an established memory.

**Figure 4.**
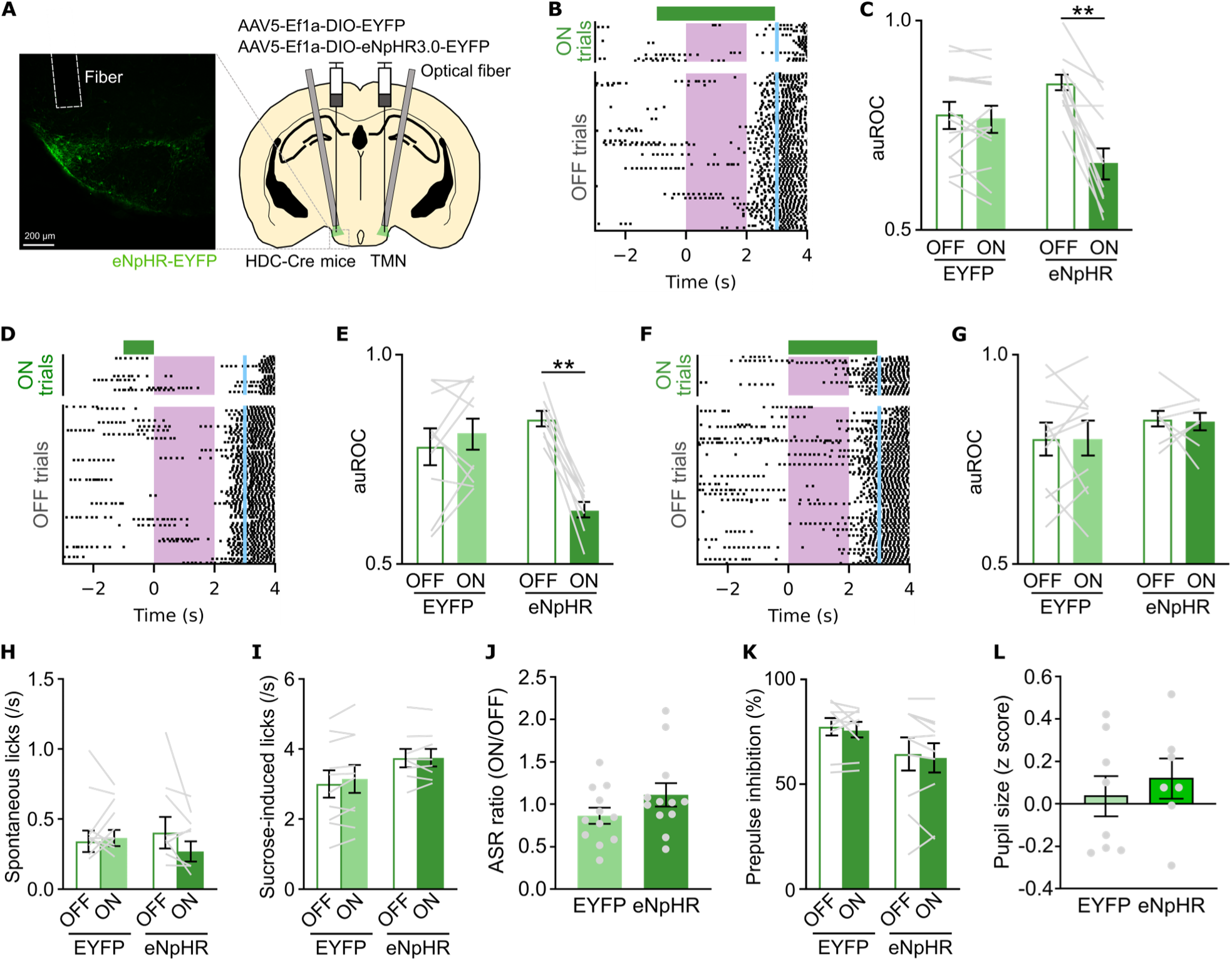
Optogenetic inhibition of histaminergic neurons attenuates memory expression. (A) Schematic of viral injections and placement of optical fibers for optogenetic inhibition of histaminergic neurons, and a representative image showing eNpHR-EYFP expression in the TMN. (B, D and F) Raster plots of licking events in a representative mouse from the eNpHR group. Green light was delivered to the TMN for 4 s (B), 1 s before CS onset (D), and 3 s after CS onset (F). (C, E and G) auROC showing changes in licking rates per trial from before to after CS, related to (B, D, F) (C: EYFP n = 13, eNpHR n = 12; E: EYFP n = 10, eNpHR n = 8; G: EYFP n = 10, eNpHR n = 8). (H–L), Inhibiting histaminergic neurons did not affect spontaneous licking (H), sucrose solution-induced licking (I), acoustic startle response (J), prepulse inhibition (K), or pupil size (L) (H, I: EYFP n = 10, eNpHR n = 8; J: n = 12; K: EYFP n = 10, eNpHR n = 13; L: EYFP n = 9, eNpHR n = 9). Data are mean ± SEM.

### Pre-CS activation of histaminergic neuronal activity enhances the accuracy of memory expression

We next tested whether pre-CS activation of histaminergic neurons is sufficient to enhance memory expression. We expressed Cre-dependent Channelrhodopsin-2 (ChR2) in TMN histaminergic neurons of HDC-IRES-Cre mice and implanted bilateral optical fibers above the TMN (Figures 5A and S6A). Blue light was delivered for 1 s (−1–0 s from CS onset). Mice were trained with two tones (CS+ paired with reward; CS– without reward), with 50 trials per tone (20 light, 30 no-light). To avoid ceiling/floor effects in performance, we analyzed the day preceding criterion performance (Figures S6B and S6C). Memory expression was quantified by auROC measuring licking discrimination between CS+ and CS–. Light delivery to the TMN increased the auROC in the ChR2 group, but not in the EYFP group (Figures 5B and 5C). Optogenetic activation did not alter spontaneous or sucrose-induced licking (Figures 5D and 5E), acoustic startle response (Figure 5F), prepulse inhibition (Figure 5G), or pupil size (Figure 5H). Thus, brief pre-CS activation of histaminergic neurons is sufficient to enhance the accuracy of memory expression.

**Figure 5.**
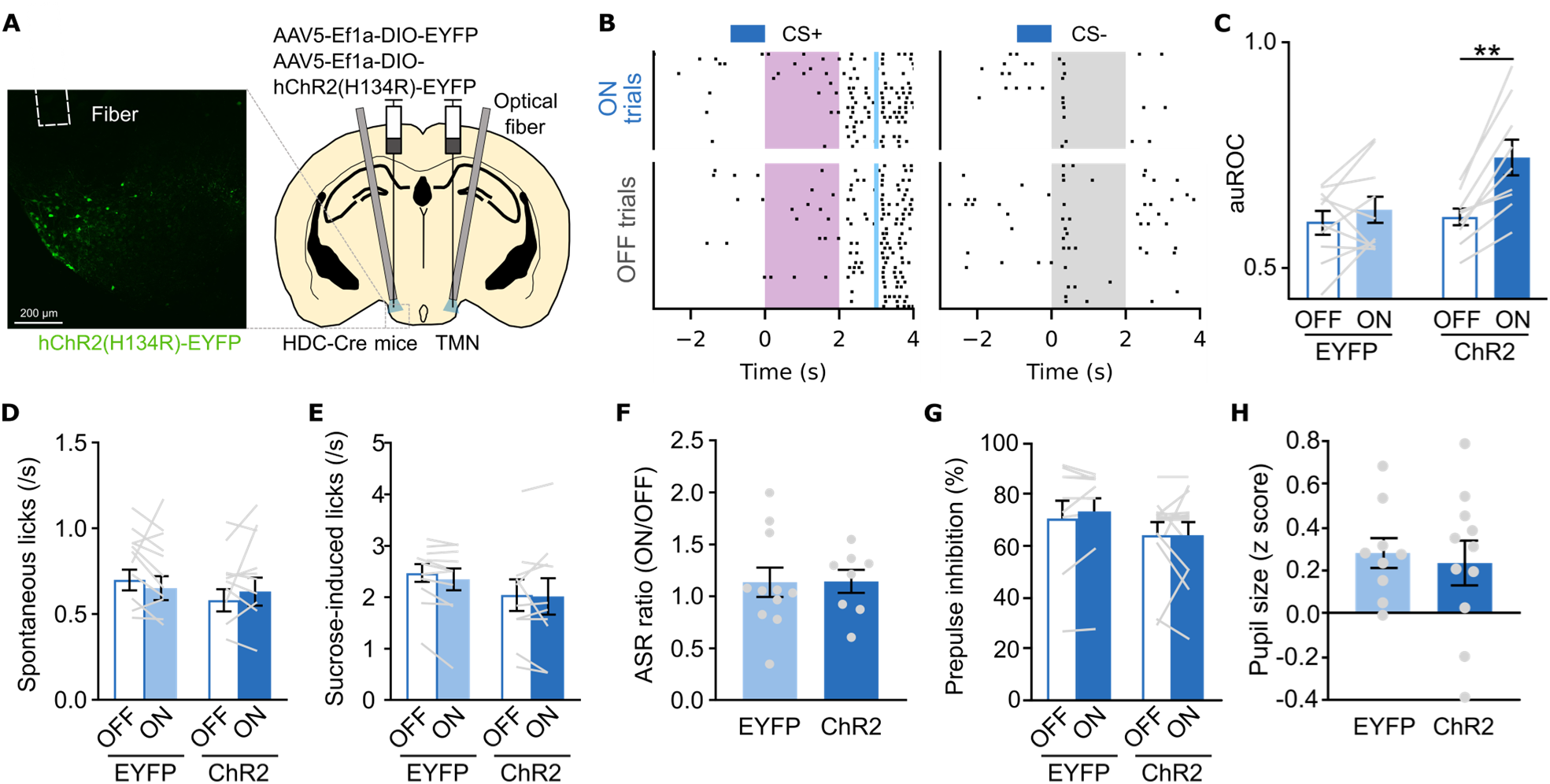
Optogenetic activation of histaminergic neurons enhances the accuracy of memory expression. (A) Schematic of viral injections and placement of optical fibers for optogenetic activation of histaminergic neurons, and a representative image showing ChR2-EYFP expression in the TMN. (B) Raster plots of licking events in response to CS+ (left) and CS− (right) from a representative mouse in the ChR2 group. Blue light was delivered to the TMN for 1 s before the onset of CS. (C) The auROC showing differences in licking rates per trial between CS+ and CS− (EYFP n = 10, ChR2 n = 9). (D–H), Activating histaminergic neurons did not affect spontaneous licking (D), sucrose solution-induced licking (E), acoustic startle response (F), prepulse inhibition (G), or pupil size (H) (D and E: EYFP n = 11, ChR2 n = 10; F: EYFP n = 11, ChR2 n = 8; G: EYFP n = 11, ChR2 n = 9; H: EYFP n = 9, ChR2 n = 11). Data are mean ± SEM.

### Direct activation of histaminergic terminals in the BLA enhances the accuracy of memory expression

The basolateral amygdala (BLA) is critical for associative learning^29,30^, and amygdala H_1_-receptor blockade impairs retrieval in inhibitory-avoidance^31^. We therefore asked whether direct activation of histaminergic terminals within the BLA is sufficient to enhance memory expression. We expressed Cre-dependent ChR2-EYFP in TMN histaminergic neurons of HDC-IRES-Cre mice and bilaterally implanted optical fibers above the BLA (Figure S6D). Post hoc histology confirmed ChR2-EYFP-positive axons in the BLA (Figure 6A). We then applied pre-CS photostimulation (−1–0 s from CS onset) during the CS+ vs. CS− discrimination task (Figure S6E). Pre-CS terminal activation increased licking discrimination (auROC) on light-ON trials relative to light-OFF trials in ChR2 mice but not in EYFP controls (Figures 6B and 6C). This terminal-specific activation did not alter spontaneous or sucrose-induced licking, acoustic startle, prepulse inhibition, or pupil size (Figures 6D–6H). Thus, brief pre-CS activation of TMN histaminergic terminals in the BLA is sufficient to enhance the accuracy of memory expression.

**Figure 6.**
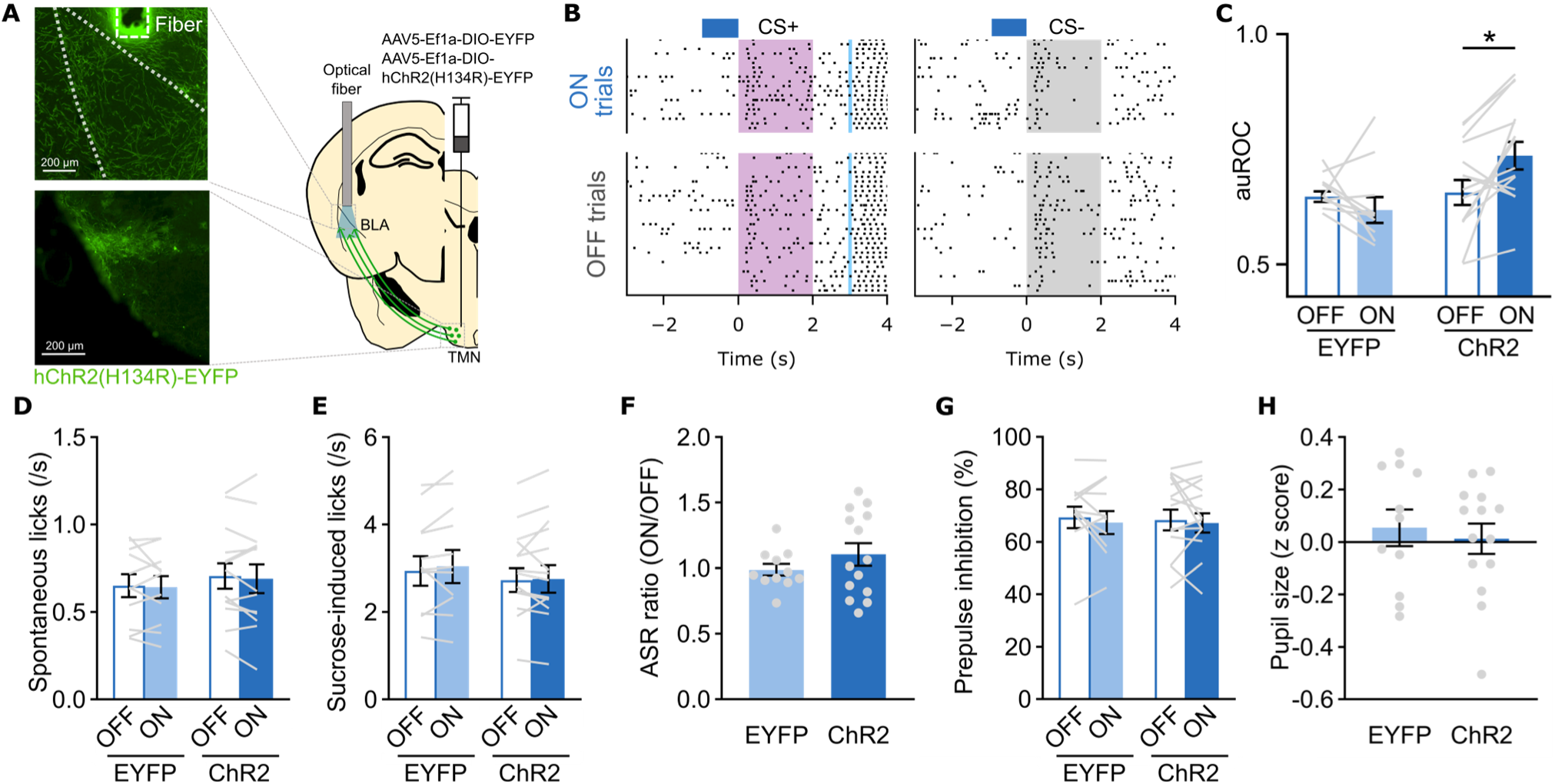
Optogenetic activation of TMN–BLA histaminergic pathway enhances the accuracy of memory expression. (A) Schematic of viral injections and placement of optical fibers for optogenetic activation of histaminergic neurons, and a representative image showing ChR2-EYFP expression in the BLA. (B) Raster plots of licking events in response to CS+ (left) and CS− (right) from a representative mouse in the ChR2 group. (C) The auROC showing differences in licking rates per trial between CS+ and CS− (n = 10 mice (EYFP), n = 14 mice (ChR2); *P = 0.0061, Sidak’s test after two-way repeated-measures ANOVA). (D–H) Activating TMN–BLA histaminergic pathway did not affect spontaneous licking (D), sucrose solution-induced licking (E), acoustic startle response (F), prepulse inhibition (G), or pupil size (H) (D–H: n = 11 mice (EYFP), n = 14 mice (ChR2)). Data are mean ± SEM.

### The CS-evoked BLA population response is amplified during high-expression trials

Our previous results identify the BLA as a key region where histaminergic input can facilitate memory expression. To understand the underlying circuit mechanisms, we next used miniaturized microscopy to record the activity of CaMKIIα-positive BLA neurons during the memory task (Figures 7A–7C and S7A). We additionally expressed eNpHR in TMN histaminergic neurons and implanted bilateral optical fibers above the TMN to permit later inhibition of histaminergic neurons (Figure 7A and S7B). First, we analyzed the relationship between BLA activity and memory expression without optogenetic manipulation (Figure S7C). After conditioning, BLA neurons exhibited heterogeneous responses to the CS, with some cells being excited, inhibited, or unresponsive (Figures 7D, 7E, and S7D). We compared CS responses between high– and non-expression trials. Population-level analysis revealed that the vector of neural responses in high-expression trials was strongly correlated with the average response vector across all trials, but with a regression slope significantly greater than 1 (Figures 7F–7H). This gain-like scaling indicates a selective amplification of the canonical (trial-averaged) CS-evoked population response during successful memory expression. In contrast, the response vector in non-expression trials showed a weaker correlation and a smaller slope compared to high-expression trials, indicating reduced alignment and gain relative to the canonical CS-evoked population response (Figures 7F–7H). At the single-neuron level, CS-excited neurons showed higher activity during high-expression trials compared to non-expression trials (Figure 7I, S7E–S7G). These differential response patterns could not be explained by licking behavior alone (Figures S7H–S7J). Together, these results indicate that successful memory expression is associated with a specific BLA population response characterized by a dynamic gain-like modulation of CS-evoked activity.

**Figure 7.**
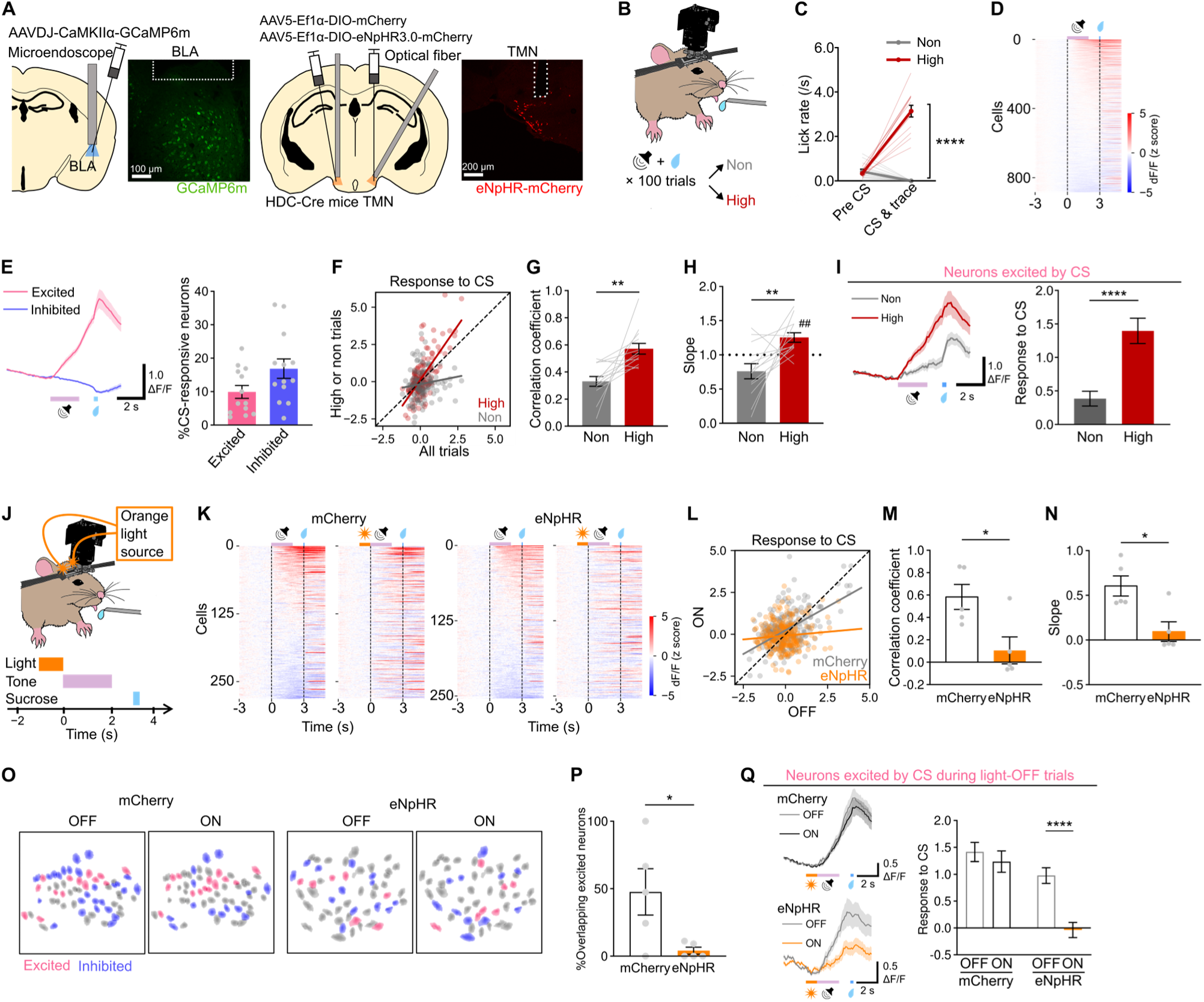
Pre-CS histaminergic neuronal activity supports the CS-evoked BLA population response. (A) Viral injections, microendoscope, and fiber placements for BLA calcium imaging with optogenetic inhibition of histaminergic neurons. (B–E) Experimental overview (B), licking rates during pre-CS and CS–trace periods (C), heatmap of neuronal activity averaged across 100 trials (D), and CS responses and percentages of CS-excited and CS-inhibited neurons (E) (n = 884 neurons, 13 mice). (F–I) CS responses of individual neurons across all trials vs. high– or non-expression trials (F), Pearson’s correlation coefficients (G), regression slopes (H), and neuronal responses of CS-excited neurons to CS (I). (J and K) TMN light delivery to inhibit histaminergic neurons before CS (J) and neuronal activity heatmaps averaged across 20 trials (K, mCherry: n = 282 neurons, 5 mice; eNpHR: n = 254 neurons, 5 mice). (L–N), CS responses of individual neurons during light-OFF vs. light-ON trials (L), Pearson’s correlation coefficients (M), and regression slopes (N). (O) Representative map of BLA neurons. (P) Proportion of neurons excited in both light conditions. (Q) CS Responses of neurons excited by CS during light-OFF trials. Data are mean ± SEM. ****P < 0.0001, **P < 0.01, *P < 0.05, ##P < 0.01 vs. chance.

### Pre-CS inhibition of TMN histaminergic neurons impairs the expression of the CS-evoked BLA population response

To test whether pre-CS histaminergic activity supports the CS-evoked BLA population response, we inhibited TMN histaminergic neurons with Cre-dependent eNpHR for 1 s (−1–0 s before CS) in the same mice while imaging CaMKIIα-positive BLA neurons (Figures 7J, 7K, and S8A). We first assessed pattern similarity and response gain by comparing neuron-wise CS response vectors between light OFF and ON trials. In the mCherry control group, the population response vectors to the CS were highly correlated between light-OFF and light-ON trials, indicating the CS-evoked population response was preserved across light conditions (Figures 7L and 7M). In the eNpHR group, the OFF–ON correlation was reduced and regression slopes were smaller than controls (Figures 7L–7N). This effect was specific to memory-related responses and did not reflect a general change in spontaneous BLA activity (Figures S8B–S8H).

We next examined the cellular basis for the reduced CS-evoked population response. In the mCherry group, 47.7% of neurons that were CS-excited in light-OFF trials were also excited in light-ON trials (Figures 7O and 7P). In the eNpHR group, however, this across-condition overlap was reduced to 4.0% (Figure 7P). The overall fraction of CS-excited neurons was unaffected by light in either group (Figure S9A and S9B). In the eNpHR group, neurons excited during light-OFF trials showed reduced CS responses during light-ON trials, whereas controls were unchanged (Figure 7Q). These effects were observed only after conditioning (Figures S9C–S9E) and were specific to the CS-excited subpopulation (Figure S9F).

Finally, we quantified population separability using a linear support vector machine trained to decode the light condition (light-OFF vs. light-ON) from trial-wise BLA population activity during the CS–trace epoch. After conditioning, cross-validated decoding accuracy in the eNpHR group exceeded chance level (50%) and was higher than in mCherry controls (Figure S9G). This indicates that pre-CS histaminergic inhibition reconfigures the CS-evoked BLA population response. Before conditioning, the decoding accuracy was comparable across groups (Figure S9H). Taken together, these findings indicate that pre-CS histaminergic neuronal activity enables the robust expression of the CS-evoked BLA population response, and that this effect is accompanied by a selective modulation of responses in CS-excited neurons.

## DISCUSSION

Moment-to-moment fluctuations in memory accessibility—epitomized by the “tip-of-the-tongue” state—are ubiquitous, yet their neurobiological origins have remained unclear. Why is a memory accessible one moment but not the next? Here, we advance a state-setting principle: spontaneous, infraslow fluctuations in histaminergic neurons create a priming state that causally gates memory accessibility. Using cell-type–specific recordings and perturbations, we show that these spontaneous dynamics, aligned with an integrated brain–body state, dynamically modulate the cue-evoked population responses in the BLA through gain-like modulation, thereby modulating memory expression. This work provides a concrete neurobiological basis for previous findings in humans that correlated pre-retrieval brain states with memory performance^5,32^. While infraslow brain activity (<0.1 Hz) has been linked to cognitive fluctuations, these studies were largely correlational and could not identify a specific cellular origin^33,34^. Here, we bridge this gap by identifying histaminergic neurons as a key cellular population whose spontaneous activity causally and dynamically modulates moment-to-moment memory accessibility. Accordingly, infraslow histaminergic dynamics may provide a temporal scaffold that enables state-dependent modulation of memory accessibility.

### Histaminergic neuronal activity and brain–body states

The fluctuations in histaminergic neuronal activity closely track a low-frequency integrated brain–body state rather than a unitary arousal signal. A cross-validated linear encoding model that combined recent histories of EEG, pupil dynamics, and facial features captured variance beyond single-modality models and reproduced the 0.05–0.1 Hz peak of the histaminergic signal, whereas no single predictor exhibited a comparable low-frequency peak. Moreover, standard arousal indices were dissociated from our effects: pupil size neither explained trial-by-trial fluctuations in memory expression nor changed with optogenetic manipulation of histaminergic neurons. Together, these observations are consistent with recent reports that TMN histaminergic neurons do not simply encode baseline arousal^35^ and suggest that their ongoing dynamics reflect an integrated, low-frequency brain–body state that modulates memory accessibility.

### A dynamic priming state model for memory accessibility

Building on our findings, we propose a dynamic priming state model that explains how histaminergic modulation is implemented at the circuit level (Figure S10). In this model, TMN histaminergic neurons act as an integrator, translating a composite brain–body state into spontaneous, infraslow fluctuations. These slow fluctuations generate a priming state, a permissive condition within memory circuits. This state does not elevate baseline activity but prepares the circuits for incoming signals. In the BLA, priming manifests as dynamic gain-like control: successful memory expression is associated not with a uniform increase in activity, but with selective amplification of the canonical CS-evoked population response. Critically, brief pre-CS inhibition of histaminergic neurons causally abolished this gain, selectively weakening responses in the CS-excited subpopulation and degrading population-code fidelity. This cue-selective reduction in effective gain, without changes in spontaneous activity, is most consistent with the loss of a priming influence. Consequently, the degree of memory expression is determined by this gain-like control: in a high-priming state, the incoming CS effectively recruits the CS-tuned ensemble and memory expression is robust; in a low-priming state, the same CS fails to drive the population response and memory expression declines. TMN histaminergic neurons can co-release GABA^36^, and the precise transmitter mechanism remains to be identified. Nevertheless, our data are more consistent with the loss of histaminergic priming over a simple disinhibition account, which would be expected to alter baseline activity.

### Timescale specialization among neuromodulators

Our work supports a specific systems-level role for histamine that complements and refines the broader neuromodulatory literature. Noradrenaline (NA) and acetylcholine (ACh) convey phasic signals related to attention, uncertainty, and plasticity, while dopamine (DA) provides phasic reinforcement– and value-related signals^37–40^; these systems also exhibit tonic state modes^41^. Within this multi-timescale landscape, ongoing histaminergic dynamics exert an infraslow, state-setting influence on memory accessibility. Together, these findings suggest functional specialization across timescales: phasic NA/ACh/DA often contribute to learning-related computations on fast timescales, whereas histaminergic dynamics provide a slower influence on the accessibility of learned information.

### Evolutionary and energetic considerations

We speculate that an infraslow histaminergic priming system provides an adaptive advantage by enabling energy-efficient cognitive readiness while dynamically tuning the exploration–exploitation balance. Sustained neural activation is energetically demanding^43,44^. Likewise, at the circuit level, maintaining a tonic high-gain state imposes a substantial metabolic cost. In addition to this general principle, prolonged histaminergic activation increases whole-body energy expenditure^42^. By contrast, the infraslow histaminergic priming we describe operates with a low duty cycle and leaves baseline activity in downstream circuits largely unchanged, providing a relatively low-cost mode of operation while keeping circuits prepared for cue-evoked responses. This infraslow fluctuation may also bias exploration–exploitation trade-offs, with low-accessibility epochs facilitating exploration and high-accessibility epochs favoring exploitation^46,47^. In this view, moment-to-moment fluctuations in memory accessibility are not neural noise but a functional feature of an energy-efficient control mode. The deep evolutionary conservation of both the histaminergic system and these cognitive fluctuations across species supports the adaptive significance of this mechanism^17,48,49^.

### Limitations of the study

Several limitations and alternative interpretations of our findings should be considered. First, while our results demonstrate that activating the TMN–BLA pathway is sufficient to enhance memory expression, we did not test its necessity by inhibiting these specific terminals. This is because interpreting terminal inhibition experiments is complex, given that rapid optogenetic silencing of axon terminals can produce paradoxical network effects or rebound^43,44^, and because directly verifying the transient suppression of spontaneous histamine release in vivo remains technically challenging^45^. Second, while our results indicate that histaminergic activity supports the CS-evoked BLA population response and that histaminergic input to the BLA can modulate memory expression, we cannot fully exclude contributions from projections to adjacent amygdalar nuclei, including the intercalated cell masses (ITCs) and the central nucleus (CeA). Anatomical studies indicate that histaminergic neurons innervate all three of these structures^14,15^. Thus, off-target light spread from optical fibers positioned above the BLA may have reached histaminergic terminals projecting to the ITCs and CeA, thereby influencing behavior. Future studies combining pathway-specific manipulation with region-specific genetic approaches could disentangle these contributions. Finally, the causal directionality among gamma-band activity, facial dynamics, and TMN output remains to be fully resolved. Simultaneous measurement of histaminergic neuronal activity, EEG, and facial movements combined with targeted perturbations would help to dissect whether these are driven by a common source or are serially coupled.

In conclusion, our findings advance histamine from a wakefulness modulator to an infraslow, state-setting modulator of moment-to-moment memory accessibility. By showing that spontaneous histaminergic dynamics establish a priming state that gates memory accessibility through gain-like modulation in the BLA, we provide a causal, circuit-level account for short-timescale fluctuations in memory accessibility. This framework explains natural fluctuation in memory accessibility and points to tractable next steps. First, non-invasive composite readouts (e.g., EEG power combined with pupil and facial indices) could be developed as biomarkers that prospectively track priming within individuals. Second, closed-loop, state-contingent interventions, rather than continuous dosing, could be evaluated to stabilize memory accessibility during identified low-priming epochs, within established safety constraints. Clarifying these predictions will be key to a systems-level understanding of cognitive dynamics in health and disease.

## STAR★Methods

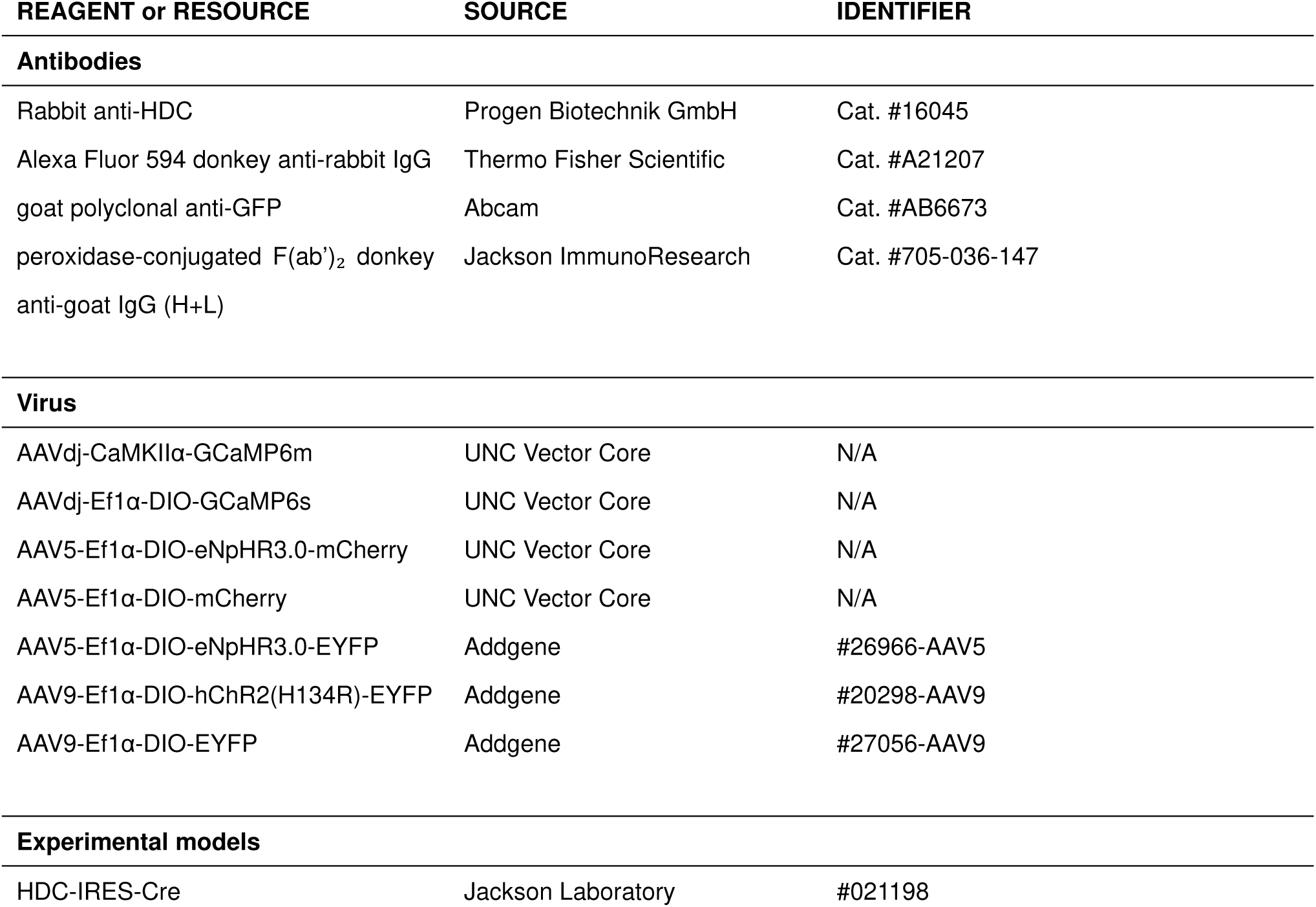
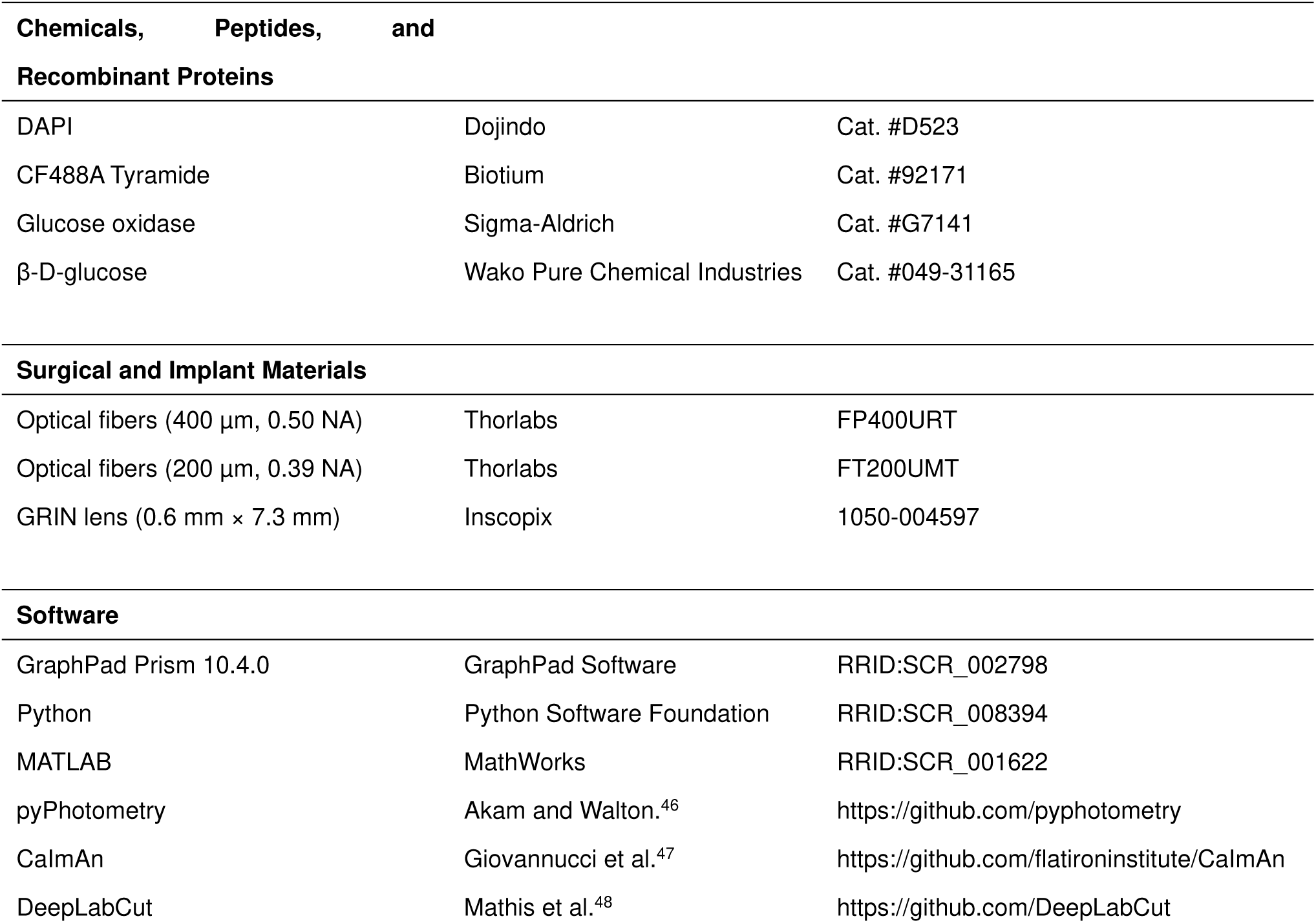
KEY RESOURCES TABLE.

## EXPERIMENTAL MODEL AND STUDY PARTICIPANT DETAILS

### Animals

Adult male and female histidine decarboxylase-IRES-Cre mice (HDC-IRES-Cre, #021198, Jackson Laboratory (Bar Harbor, Maine, USA))^49^, aged 7–15 weeks, were housed individually. They were maintained on a 12-hr light–dark cycle, with lights on at 07:00 AM, and had unrestricted access to food and water unless otherwise noted. Behavior tests were conducted during the light phase of this cycle. All animal experiments were approved by the Institutional Animal Care and Use Committee of Hokkaido University (approval number: 16-0043) and Nagoya City University (approval number: 22-018). The study adhered to the Hokkaido University and Nagoya City University guidelines for the care and use of laboratory animals and complied with national guidelines, including the Guidelines for Proper Conduct of Animal Experiments (Science Council of Japan), the Fundamental Guidelines for Proper Conduct of Animal Experiments and Related Activities in Academic Research Institutions (Ministry of Education, Culture, Sports, Science and Technology, Notice No. 71 of 2006) and, the Standards for Breeding and Housing of and Pain Alleviation for Experimental Animals (Ministry of the Environment, Notice No. 88 of 2006).

## METHOD DETAILS

### Surgery

The viral vectors AAVdj-CaMKIIα-GCaMP6m (UNC Vector Core (Chapel Hill, North Carolina, USA)), AAVdj-Ef1α-DIO-GCaMP6s (UNC Vector Core), AAV5-Ef1α-DIO-eNpHR3.0-mCherry (UNC Vector Core), AAV5-Ef1α-DIO-mCherry (UNC Vector Core), AAV5-Ef1α-DIO-eNpHR3.0-EYFP^50^ (Addgene (Watertown, Massachusetts, USA) viral prep #26966-AAV5), AAV9-Ef1α-DIO-hChR2(H134R)-EYFP (Addgene viral prep #20298-AAV9), and AAV9-Ef1α-DIO-EYFP (Addgene viral prep #27056-AAV9), kindly provided by Karl Deisseroth, were used in the study.

Before surgery, the mice were intraperitoneally injected with carprofen (5 mg/kg) and dexamethasone (0.2 mg/kg). They were then anesthetized with isoflurane (0.8–1.5%), and positioned on a stereotaxic apparatus (SR-6M-HT, Narishige, Tokyo, Japan) kept warm with a heating pad. Lidocaine (2%; Aspen Japan, Tokyo, Japan) was applied to the scalp to relieve pain. The scalp was cut along the centerline and small holes were drilled above the injection sites. For fiber photometry experiments, we injected AAVdj-Ef1α-DIO-GCaMP6s (3.9×10^12^ vg/mL) into the bilateral TMN (A/P: −2.5 mm, M/L: ±0.92 mm, D/V: −5.4/5.45 mm) at a rate of 0.1 µl/min. Following the virus injection, optical fibers (400 µm, 0.50 NA; Thorlabs, Newton, New Jersey, USA) with stainless-steel ferrules were implanted 300 µm above the virus injection sites and were secured to the skull with a self-curing adhesive resin cement (Super-Bond, SUN MEDICAL, Moriyama, Japan) and black dental cement (#1530BLK; Lang Dental, Wheeling, Illinois, USA). Additionally, a chamber frame (CF-10, Narishige, Tokyo, Japan) was attached to the skull with dental cement (UNIFAST III; GC, Tokyo, Japan) for behavioral experiments, with the head fixed. After surgery, carprofen, dexamethasone, and amoxicillin (50 mg/kg) were administered for 7 days.

For simultaneous EEG and fiber photometry recordings, three stainless-steel pan-head screws (M1.0 × 2 mm length; ESCO, Osaka, Japan) were implanted unilaterally into the skull after implanting an optical fiber. All three screws were placed ipsilaterally (on the same side of the brain). A recording screw was placed over the medial prefrontal cortex (A/P: +1.5 mm) and another over the anterior hippocampus (A/P: −1.0 mm), with a reference screw anchored over the cerebellum (A/P: −6.0 mm). The medial-lateral coordinate for all screws was identical (either M/L: +1.5 mm or −1.5 mm).

For optogenetic manipulation of histaminergic neurons, a virus encoding the Cre-inducible eNpHR-EYFP, ChR2-EYFP, or EYFP (AAV5-Ef1α-DIO-eNpHR3.0-EYFP (1.0×10^13^ vg/mL), AAV9-Ef1α-DIO-hChR2(H134R)-EYFP (1.0×10^13^ vg/mL), or AAV9-Ef1α-DIO-EYFP (1.0×10^13^ vg/mL)) was injected into the bilateral TMN (A/P: −2.5 mm, M/L: ±1.0 mm, D/V: −5.45 mm) of HDC-IRES-Cre mice. After the virus injections, optical fibers (200 µm, 0.39 NA, Thorlabs) with ceramic ferrules were implanted 300 µm above the virus injection sites of TMN. The fibers were implanted at a 10° angle on the midline through the hole targeting the TMN (A/P: −2.5 mm, M/L: ±1.9 mm, D/V: −5.0 mm). For photostimulation of the TMN histaminergic terminals in the BLA, optical fibers were implanted 300 µm above the BLA (A/P: −1.4 mm, M/L: ±3.0 mm, D/V: −4.2 mm). The fibers and chamber frame positioned above the skull were secured to the skull using self-curing adhesive resin cement and black dental cement.

Two AAV vectors were injected for Ca^2+^ imaging using optogenetic manipulation. First, AAVdj-CaMKⅡa-GCaMP6m (1.9×10^13^ vg/mL) was injected into the unilateral BLA (A/P: −1.7 mm, M/L: ±3.35 mm, D/V: −4.7 mm). We also injected either AAV5-Ef1α-DIO-eNpHR3.0-mCherry (4.0×10^12^ vg/mL) or AAV5-Ef1α-DIO-mCherry (5.1×10^12^ vg/mL) into the bilateral TMN (A/P: −2.5 mm, M/L: ±1.0 mm, D/V: −5.45 mm). After the virus injections, optical fibers (200 µm, 0.39 NA, Thorlabs) with ceramic ferrules were implanted 300 µm above the virus injection sites of TMN. Subsequently, a Gradient Index (GRIN) lens (0.6 mm in diameter, 7.3 mm in length; Inscopix, Palo Alto, California, USA) was implanted 100 µm above the BLA injection site. Both the fibers and GRIN lens were secured to the skull using a self-curing adhesive resin and black dental cement. The GRIN lens was covered with Kwik-Sil (World Precision Instruments, Sarasota, Florida, USA). The injections and fiber implantations were performed at a 30° lateral angle for one TMN and a 30° posterior angle for the other.

Baseplate surgery was performed more than 5 weeks after implantation of the GRIN lens. The mice were anesthetized with isoflurane (0.8–1.5%), and placed on the stereotaxic apparatus kept warm with a heating pad. After removing the Kwik-Sil, a miniscope (UCLA miniscope V4, Open Ephys, Lisbon, Portugal) with a baseplate was positioned to optimally view the blood vessels and GCaMP6m dynamics. The baseplate was secured to the previously formed cement by using additional cement. After the baseplate was attached, the miniscope was detached, and a plastic cover was placed on it to protect the GRIN lens.

### Tissue processing

Following the behavioral experiments, the mice were deeply anesthetized using a combination of medetomidine (0.3 mg/kg; ZENOAQ, Fukushima, Japan), midazolam (4.0 mg/kg; Sandoz K.K., Tokyo, Japan), and butorphanol (5.0 mg/kg; Meiji Animal Health Co., Ltd., Kumamoto, Japan). They were then transcardially perfused with phosphate-buffered saline (PBS) (30 mL), followed by 4% paraformaldehyde (PFA) in 0.1 M phosphate buffer (PB) (30 mL). The brains were removed and further fixed in 4% PFA overnight at 4°C, followed by cryopreservation for 48–72 h in 15% and 30% sucrose solutions in PB at 4°C before freezing. Coronal brain sections, each 40 μm thick, were prepared using a freezing microtome (CM1900 or CM3050S, Leica, Wetzlar, Germany). The sections were mounted onto glass slides with a mounting medium (20 mM Tris, 0.5% N-propyl gallate, 50–90% glycerol, pH 8.0). Images were acquired using a fluorescence microscope (Nanozoomer S60, Hamamatsu Photonics, Hamamatsu, Japan; BZ-X700, Keyence, Osaka, Japan; A1RS+, Nikon, Tokyo, Japan; or FV3000, Evident, Tokyo, Japan) to verify the exact placement of the fiber or lens implants and the expression of GCaMP6s, GCaMP6m, eNpHR3.0-EYFP, eNpHR-mCherry, ChR2-EYFP, EYFP, and mCherry.

### Immunohistochemistry

Immunohistochemistry was performed after tissue processing. First, sections were incubated in PBST (0.1% Triton X-100 in PBS) for 15 min. They were then blocked in PBS-BX (0.01 M PBS/3% BSA, 10.25% TritonX-100) for 1 h at RT. The sections were then incubated with rabbit polyclonal antibodies against HDC (1:800, Cat. #16045; Progen Biotechnik GmbH, Heidelberg, Germany) at 4°C. After overnight incubation, the sections were washed three times with PBS-BX (15 min each) and incubated with Alexa Fluor 594 donkey anti-rabbit IgG (1:1000, Cat. #A21207; Thermo Fisher Scientific, Waltham, Massachusetts, USA) for 2 h at RT, followed by 5-min staining with DAPI (0.3 µg/ml, 4′,6-diamidino-2-phenylindole, Dojindo, Cat. #D523) in PBS. After two washes with PBS (5 min each), sections were mounted onto glass slides with a mounting medium (20 mM Tris, 0.5% N-propyl gallate, 50–90% glycerol, pH 8.0). To visualize EYFP-labeled histaminergic terminals, a tyramide signal amplification protocol^51^ was employed. Sections were incubated in 1% H₂O₂ in PBS for 30 min to inactivate endogenous peroxidase, washed twice in 0.3% PBS-X (0.3% Triton X-100 in PBS; 10 min each), and incubated overnight with goat polyclonal anti-GFP (1:5000; Abcam, Cat. #AB6673) diluted in PBS-XCD (PBS-X containing 0.12% λ-carrageenan, 1% normal donkey serum, and 0.02% thimerosal). After two washes, sections were incubated for 4 h with peroxidase-conjugated F(ab’)₂ donkey anti-goat IgG (H+L) (1:500; Jackson ImmunoResearch) diluted in PBS-XCD. The sections were then washed twice, rinsed twice with 0.1 M phosphate buffer (5 min each), and incubated for 10 min in CF488 FT-GO solution (10 µM CF488A tyramide (Biotium, Cat. #92171) and 3 µg/mL glucose oxidase in 2% BSA/0.1 M phosphate buffer). The FT-GO reaction was initiated by adding β-D-glucose solution (2 mg/mL in distilled water) and continued for 20 min. Sections were then washed twice with 0.3% PBS-X, rinsed in PBS, and mounted with 50% glycerol in PBS. Images were obtained using a laser scanning confocal microscope (A1RS+, Nikon or FLUOVIEW FV10i, Evident).

### Tone-reward conditioning with fiber photometry

We used a custom-made behavioral apparatus equipped with a licking spout, an infrared sensor to detect licking, and speakers. The entire system was managed using Python-based custom software running on a single-board computer (Raspberry Pi 4 B). Mice were food-restricted to maintain their weight at 80–85% of the weight they had with free access to food. Initially, each mouse was habituated to being head-fixed and trained to lick a spout dispensing 10% sucrose solution (3 µl) for 2–8 days through a gravity-driven, solenoid-controlled lick tube. During this habituation phase, the sucrose solution was presented 100 times at pseudorandom intervals between 5 and 11 s. Once the mice reliably licked the spout (at > 1,000 licks per 100 sucrose presentations), they were habituated to a 2-s tone (conditioned stimulus, CS, either 4 kHz, 7 kHz, 10 kHz, 65 dB) 50 times, with the tone frequency varying depending on the mouse.

The following day, the mice were subjected to tone-reward conditioning. Each trial began with the 2-s CS, followed by a 1-s delay, and then a 10% sucrose reward (3 µl, unconditioned stimulus, US). Conditioning consisted of 100 trials per day for 3–14 days. The interval between trials varied randomly between 23 and 29 s. Fiber photometry recording was performed on the first day of tone-reward conditioning and the day after the mice learned the association between the CS and US (evidenced by a significant increase in licks for 3 s after tone onset compared to 3 s before, P < 0.01, Wilcoxon signed-rank test). Although the licking rate does not always precisely reflect the degree of memory expression, we can make certain inferences: if the licking rate is zero, it suggests that memory expression has minimally occurred; if the rate falls within the top 10% of all trials, we can consider the degree of memory expression to be high. Therefore, trials in which the licking rate for 3 s after the tone onset was in the top 10% were categorized as high-expression trials. Of the trials in which the mice did not lick, the first 10% of trials were classified as non-expression trials. This is because the absence of licking in later trials could be due to satiation of the mice rather than a failure to express the association. Mice with fewer than five non-expression trials were excluded from the analysis.

### Conditioning with optogenetic inhibition

For tone-reward conditioning with optogenetic inhibition, we employed a procedure similar to that used in our fiber photometry experiments, with the following modifications. Once we confirmed that the mice had reliably licked the spout, we initiated conditioning sessions instead of tone habituation. We used a 4-kHz, 75-dB tone as the CS. The mice underwent 100 conditioning trials a day for 3–14 days (EYFP group: 10.4±0.7 days, eNpHR group: 9.0±1.2 days) until they learned the association between the CS and US. After the expression performance met our criteria (evidenced by a significant increase in licks for 3 s after tone onset compared to 3 s before tone onset, P < 0.01, Wilcoxon signed-rank test), optogenetic experiments were initiated. On these days, mice underwent 20 light-ON trials and 80 light-OFF trials in a pseudorandom order. We delivered constant green light (10 mW) to the bilateral TMN using an LD-pumped all-solid-state laser (MGL-III-532 nm-300 mW, CNI laser, Changchun, China) for 4 s from 1 s before CS onset until sucrose presentation, for 1 s before CS onset, or for 3 s from CS onset until sucrose presentation. We calculated the auROC to assess the difference in licking rates per trial for 3 seconds before and after CS onset.

To assess whether optogenetic inhibition directly affected licking behavior, we intermittently presented sucrose solutions without CS at pseudorandom intervals between 20 and 32 s across 100 trials. Green light was delivered to the TMN for 9 s starting 6 s before the onset of sucrose presentation during the 20 trials. We evaluated spontaneous and sucrose-induced licking by analyzing licking rates during the 6 s before and the 3 s after the onset of sucrose presentation.

### Conditioning with optogenetic activation

Once we confirmed that the mice reliably licked the spout, we performed conditioning sessions using two 2-s CS (7 kHz and white noise, both at 75 dB) with daily light delivery. Each session included 50 trials of both CS+ and CS−. The CS+ was followed by a 1-s delay and a sucrose solution reward, while CS− was not rewarded. We delivered a blue light (5 mW) to either the bilateral TMN or BLA using a DPSS laser (MBL-III-473-100mW, CNI laser) during 20 trials each of CS+ and CS− in a pseudorandom order. The laser was turned on for 5-ms pulses (10 Hz) for 1 s before CS onset. Each day, we assessed learning completion based on the performance in light-OFF trials for each mouse. Learning was considered complete if the licking rates for 3 s after the CS+ onset were significantly higher than both the rates for 3 s before the CS+ onset and after the CS− onset (P < 0.01; Wilcoxon signed-rank test and Mann–Whitney U test, respectively). Memory performance during light-ON versus light-OFF trials was compared using data from the day before learning was completed. We calculated the auROC to assess the difference in licking rates per trial for 3 s following the CS+ and CS− onset. Experiments evaluating whether optogenetic activation affects licking behavior were conducted similarly to those with optogenetic inhibition, except that we used 10 Hz blue light delivery.

### Fiber photometry

Fiber photometry experiments were performed at a minimum of 3 weeks after virus injection. Except for the bilateral recordings, the recordings were performed using either an sCMOS camera or a photomultiplier for data collection. A previously established protocol was used for the sCMOS camera^52^. Briefly, fluorescence was captured using a patchcord comprised of four 400-μm-diameter fibers with an NA of 0.48 (Thorlabs). This patch cord was connected to an SMA connector positioned at the working distance of the objective, and images were captured using a 20x/0.75 NA objective. Images were recorded using a sCMOS camera (ORCA-Flash4.0 V3; Hamamatsu Photonics). The patch cord was terminated using stainless-steel ferrules. One of these ferrules was connected to another ferrule implanted in the mouse via a ceramic sleeve. To monitor GCaMP6s activity, we used 405-nm and 470-nm LEDs (M470F3, M405F1; Thorlabs). The LEDs were fiber-coupled, merged using a 425-nm long-pass dichroic mirror, and integrated into the system with a 495-nm long-pass dichroic mirror. Signal collection and analysis were performed using a custom MATLAB script (45). The recordings were conducted at 15 Hz with alternating frames of 470 nm and 405 nm excitation wavelengths, producing a frame rate of 7.5 Hz each for both GCaMP6s Ca^2+^ and isosbestic control signals. For photomultiplier-based recordings, a photometry system with integrated 470 nm and 405 nm LEDs (ilFMC4-G1_IE(400-410)_E(460-490)_F(500-550)_S, Doric, Quebec, Canada) was used. The LEDs were combined via a dichroic mirror in the minicube and connected to the optical fiber patch cable (400 μm, 0.57 NA, Doric). The GCaMP6s emission was detected using a photomultiplier (C7169, Hamamatsu Photonics) and digitized using pyPhotometry^46^. The GCaMP6s Ca^2+^ and isosbestic control signals were acquired at 130 Hz. The LED intensity at the tip of the patch cable (60–90 μW) was kept constant for each mouse across the sessions.

For recordings from the bilateral TMN, images acquired via optical fibers implanted into the brain were captured using a macro zoom microscope (MVX10; ×2 objective, 0.50 NA; Evident). GCaMP6s fluorescence was excited with blue light (470 nm LED; Thorlabs), and the resulting fluorescence was acquired at a resolution of 512 × 512 pixels using a high-speed CMOS camera (ORCA-Flash4.0) at a frame rate of 33.3 fps.

### Simultaneous recordings of fiber photometry, EEG, and facial dynamics

Mice were habituated to head fixation in the custom experimental chamber for 20 min per day on 3 consecutive days. On the recording day, mice were head-fixed, and after a 2-min stabilization period, spontaneous activity was recorded for 20 min. Fiber photometry was used to unilaterally record GCaMP fluorescence signals from histaminergic neurons. Cortical EEG signals were amplified (HAS-4; Bio Research Center, Nagoya, Japan) and digitized at 200 Hz using a USB-6002 data-acquisition device (National Instruments). Facial movements, including those of the eye, nose, mouth, ear, whiskers, and pupil, were captured at 120 frames per second using a no-IR camera (RPI-SC0873, Raspberry Pi Ltd., Cambridge, UK) controlled by a custom Python script on a Raspberry Pi 5. The face was illuminated with infrared LED ring light (FRS5CS, 850 nm, OptoSupply). To synchronize all data streams, the Raspberry Pi generated TTL pulses, which were simultaneously sent to the USB-6002 and the fiber-photometry acquisition system (pyPhotometry).

### Closed-loop CS–US presentations

Closed-loop CS–US presentations were performed based on the activity of histaminergic neurons. The Windows PC for recording photometry signals was equipped with a pyPhotometry acquisition board and an I/O board (Arduino Uno) to send the TTL to the Raspberry Pi. Custom-made Python-written software, modified from the pyPhotometry acquisition software, detected the dynamics of the photometry signal and sent TTL signals to the Raspberry Pi to output the CS and US. The 470-nm and 405-nm signals were each recorded at 130 Hz, but only the raw 470-nm signal was used to determine the CS–US presentations. To stabilize the photometry signal, we kept the mice head-fixed for 12 min. The signal measured between 10–12 min was used as baseline. Subsequently, CS–US was performed when the histaminergic neuronal activity was either high or consistently low. The order of presentation of the CS–US during high or low histaminergic neuronal activity was predetermined randomly. The session lasted for 60 min. The CS–US was presented when all the following conditions were met: a minimum of 25 s had passed since the last CS–US presentation, the mouse had not licked in the preceding 3 s, and the median of the three most recent signal samples exceeded an upper threshold or did not exceed a lower threshold for 10 s. The thresholds were determined as follows:

Upper threshold = median of the last 20-s signals + (the value of the top 2% of baseline signals minus the median of baseline signals).

Lower threshold = median of the last 20-s signals + (the value of the top 10% of baseline signals minus the median of baseline signals).

### *In vivo* calcium imaging

Calcium imaging was performed using a UCLA Miniscope V4^53^. For optogenetic manipulation, orange light stimulation (10 mW) was delivered using a laser diode-based light source (LISER_LED470, Doric) and passed through a bandpass filter (LBPF_612/069, Doric). To synchronize imaging and optogenetic manipulation, imaging timestamps were recorded on a Raspberry Pi, which also transmitted TTL pulses to a light source. Prior to imaging, mice underwent habituation sessions for several days. In each habituation session, they were head-fixed and a dummy miniscope was attached to the baseplate. The mice were trained to lick a spout dispensing 10% sucrose solution, which was presented 100 times at random intervals ranging between 5 and 11 s. Once the mice showed consistent licking behavior, defined as > 1,000 licks per 100 sucrose presentations, they underwent a series of imaging sessions. Initially, we observed the effect of inhibiting histaminergic neuronal activity on baseline BLA activity. Light stimulation (10 trials) and control trials without stimulation (10 trials) were conducted alternately. In each light stimulation trial, a 1-s orange light stimulus was delivered to the TMN. The intertrial intervals were randomized between 13 and 19 s. We then recorded the BLA activity in response to 40 trials of a 2-s tone (CS, 4 kHz, 75 dB). In half of these trials, a 1-s orange light was delivered to the TMN 1 s before the CS onset. After the third day, mice underwent tone-reward conditioning. After mice learned the CS–US association, indicated by a significant increase in licks during the CS period compared to the pre-CS period (both 3 s in duration; P < 0.01, Wilcoxon signed-rank test), we imaged BLA neuronal activity across 100 trials without light delivery. The high-expression and non-expression trials were defined in the same manner as in the photometry experiments. The next day, activity was imaged during 40 tone-reward conditioning trials, with a 1-s orange light delivered to the TMN before half of the tone onsets. Throughout conditioning, the intertrial intervals varied randomly between 13 and 19 s.

### Acoustic startle response

The acoustic startle response test was performed on head-fixed mice. The behavioral apparatus was based on a startle response system equipped with a speaker and motion sensor (SR-LAB, San Diego Instruments, Inc., San Diego, California, USA). In our modified setup, the analog output indicating the startle response was recorded by a Raspberry Pi, which provided analog input for sound presentation. To synchronize the measurement of the startle response with optogenetic manipulation, the same Raspberry Pi controlled the green and blue lasers. Before the test session, background noise (60 dB) was emitted from the speaker for 300 seconds to acclimatize the mice. The test session consisted of 10 trials of startle stimuli (90 dB, 100 ms), and 5 out of 10 trials involved light stimulation of either the bilateral TMN or BLA for 1.1 s starting from 1 s before the startling pulse. These pulses were presented at four different intertrial intervals: 12, 16, 20, and 24 s. The maximum signal following the onset of the startling pulse was used for the analysis.

### Prepulse inhibition

The prepulse inhibition (PPI) test was performed on head-fixed mice using the same apparatus used for the acoustic startle test. The PPI test consisted of four different conditions, with 10 trials per condition: (1) Pulse alone: a single 120-dB startling pulse (40 ms) to establish baseline startle amplitude; (2) Prepulse + pulse: a 75-dB prepulse (20 ms) followed by the startling pulse after 100 ms; (3) Light stimulation + pulse: a 1.1-s light stimulation preceding the startling pulse, without prepulse; and (4) Light stimulation + prepulse + pulse: a 1.1-s light stimulation period ending at the startling pulse onset, with the prepulse delivered 100 ms before the startling pulse. These trials were presented at four different intertrial intervals: 16, 19, 22, and 25 s. The maximum signal following the onset of the starting pulse was used for the analysis. Green or blue light was delivered bilaterally to the TMN or BLA.

### Pupillometry

Videos of one eye of each mouse were captured using a no-IR Raspberry Pi Camera Module V3 (RPI-SC0873, Raspberry Pi Ltd., Cambridge, UK) at a frame rate of 30 fps. An infrared light (780 nm; LIU780A, Thorlabs) was used to illuminate the eye. To examine the relationship between histaminergic neuronal activity and pupil size, we performed pupillometry with fiber photometry for 10 min. In the optogenetic manipulation experiments, we measured pupil size across 20 trials; during 10 trials, green or blue light was delivered to the TMN for 4 s. For pupil tracking, we employed DeepLabCut (ver. 2.2rc3)^48^. Eight points around the pupil were manually annotated, and this labeled dataset was used to train the model using a ResNet-50-based neural network. After verifying the accuracy of the label positioning, the software automatically determined the positions of the eight points in the other video frames from all mice. The pupil size was computed using a custom Python script based on the acquired pupil location data. An ellipse was fitted to an octagon, defined by the label coordinates, to determine pupil size, which was normalized to the z-score. The temporal derivative of the pupil size signal was calculated to capture dynamic changes in pupil size. Using Numpy^54^, the gradient of the pupil size with respect to time was computed to yield derivative values representing the rate of change in pupil size. A smoothing step was applied to the derivative by using a Hann window (30-frame size, 1 s) from SciPy^55^ to reduce noise. For optogenetic inhibition (Figure 4L), green light was delivered to the TMN for 4 s, and the mean pupil size during the latter 3 s was compared. For optogenetic stimulation, blue light was delivered to the TMN (Figure 5H) or BLA (Figure 6H) for 1 s, and the mean pupil size during the following 3 s was compared.

### Data analysis for fiber photometry

The photometric data from the photomultiplier were processed as follows. Both the 470-nm and 405-nm signals were median-filtered (kernel size of 5) and low-pass filtered (10 Hz). An exponential fit was applied to the filtered signals to eliminate slow bleaching. The normalized change in fluorescence (dF/F) was calculated by subtracting the median of the entire signal and dividing it by the median value. To correct for motion in the 470-nm signal, we scaled the 405-nm reference dF/F signal to best fit the 470-nm dF/F signal using least-squares regression, and subtracted this from the 470-nm signal. For baseline determination, we pooled data from the 10-s pre-trial period to trial onset. This approach was applied across all the trials to calculate a single mean and standard deviation for each session’s baseline. The motion-corrected signal was normalized by subtracting the mean and dividing by the baseline standard deviation. The sCMOS camera data analysis followed a similar protocol, except that double exponential fitting was used to effectively remove slow bleaching instead of median filtering, low-pass filtering, and exponential fitting. To obtain histaminergic neuronal activity immediately before CS onset, we extracted the signal from a single time point, specifically two time points (267 ms) before CS onset, to prevent contamination from neuronal activity induced by CS presentation. The median signal values were calculated separately for high-expression and non-expression trials and used for comparisons between the two conditions.

### Analysis of spontaneous histaminergic neuronal activity, EEG, and facial movements

EEG signals were resampled to 10 Hz. EEG was bandpass-filtered into very-low-frequency (0.01–0.5 Hz), delta (0.5–4 Hz), theta (4–8 Hz), alpha (8–12 Hz), beta (12–30 Hz), and gamma (30–80 Hz), and the Hilbert envelope was extracted.

The two-dimensional coordinates of multiple facial landmarks were tracked using DeepLabCut. These included the eye (4 points), nose (3 points), whisker pad (3 points), mouth/jaw (2 points), and ear (3 points), as well as 8 points delineating the pupil. All landmark trajectories were interpolated to 10 Hz. For non-pupil landmarks, principal component analysis (PCA) was performed. This analysis was performed independently for each facial region and for each mouse, and the first three principal components (PCs) were retained as features. For the pupil, its size was computed from the 8 tracked points described above.

For power spectral density (PSD) analysis, all signals were resampled to a uniform sampling rate of 10 Hz and z-scored. The PSD was then estimated using the Welch’s method from the SciPy library with a frequency resolution of 0.01 Hz. The resulting PSD profile was normalized by its total power within the 0–0.5 Hz frequency range. For EEG, PSDs were calculated for each of the two channels and then averaged prior to normalization. For facial movements, the PSDs of the top three principal components from each region were summed before the normalization step.

To examine the temporal relationships between histaminergic activity and other physiological signals, a cross-correlation analysis was performed. Facial movement was quantified as motion energy derived from the first three principal components of the tracked facial landmarks. All signals were low-pass filtered at 1 Hz and normalized to their respective z-scores. Cross-correlation functions were then computed using the signal.correlate function in the SciPy library. For EEG signals, the cross-correlation functions were computed for each channel separately and then averaged across channels.

### Encoding model to predict histaminergic neuronal activity

A linear encoding model was constructed using Ridge regression to predict the GCaMP signal from EEG, pupil, and facial movement features. All data were resampled to 10 Hz. GCaMP signal was band-pass filtered (0.01–1.0 Hz). EEG features were obtained by filtering each channel into six bands (very-low-frequency–gamma) and extracting the Hilbert envelope. In addition, we used causal 10-s backward moving averages of these envelopes. Pupil features included the pupil size and its temporal derivative. Facial movement features were obtained by applying PCA separately to each facial part and retaining the top three PCs. The design matrix included a 2-s history of all predictors, using only past lags.

Model performance was assessed using a nested time-series cross-validation scheme implemented with TimeSeriesSplit in scikit-learn, with 3 outer folds and 3 inner folds. To prevent temporal leakage across fold boundaries, we enforced a 2-s gap between the last training sample and the first test sample within each split. The inner loop optimized the Ridge regularization parameter (alpha) using only the training data, and the outer loop provided an unbiased estimate of generalization by scoring the held-out test segments. The model’s generalization performance was reported as cvR², defined as the coefficient of determination (R²) averaged across the outer test folds.

The significance of the group-mean cvR² was assessed using a permutation test. For each mouse, a null distribution was generated by recomputing the cvR² 1,000 times with the GCaMP signal circularly shifted by a random interval of at least 30 s. A group-level null distribution was then constructed by repeatedly (100,000 iterations) calculating the mean of values randomly sampled from each mouse’s null set. The one-sided p-value was defined as the proportion of null group means greater than or equal to the observed group-mean cvR². To evaluate the contribution of individual feature groups, we performed two additional analyses. First, we quantified the predictive power of each group in isolation by training models on single feature groups and evaluating their cvR². Second, we measured the importance of each group to the full model by calculating the drop in performance (ΔcvR²) after disrupting the group’s contribution. This disruption was achieved by circularly shifting the block of time series corresponding to the feature group on the held-out test data. This procedure was repeated 100 times with random shift lengths, and the mean ΔcvR² was reported. For both analyses, a ‘feature group’ was defined as a primary feature and all its associated derivatives. For example, the ‘pupil’ group included both pupil size and its temporal derivative, the ‘gamma’ group included the gamma-band power from both EEG channels plus their 10-s moving averages, and the ‘nose’ group included all three of its PCs.

### Data analysis for calcium imaging

Calcium imaging data were analyzed using the CaImAn software^47^ according to our previous study (39). Initially, raw imaging data were subjected to motion correction. Subsequently, regions of interest (ROIs) and the time series of their corresponding fluorescence signals were generated using constrained non-negative matrix factorization. To ensure accuracy, automatically extracted ROIs were subjected to manual quality control checks. When the outer fiber tract surrounding the BLA was visible in the field of view, we excluded ROIs from the endopiriform nucleus side and analyzed only those within the BLA (8). For each ROI, we calculated dF/F, which represented the change in fluorescence relative to the baseline fluorescence level. The dF/F traces were deconvolved to infer neuronal activity (spiking activity). The deconvolved activity (arrays of estimates S generated by CaImAn) was estimated to be approximately proportional to the firing rate.

To assess individual neuronal responses to CS, the sum of deconvolved activity per second was calculated for a 3-s window before and after CS onset. In calcium imaging across 100 trials without light delivery, neurons were classified as excited or inhibited by CS if their deconvolved activity following CS onset was significantly higher or lower, respectively, than their pre-CS baseline activity (Wilcoxon signed-rank test, P < 0.05). For neuronal classification in imaging sessions with light delivery, we calculated the auROC for each neuron by comparing responses following CS presentation to the baseline activity before CS onset during light-OFF or –ON trials. Neurons with auROC values exceeding the threshold (0.576) were classified as excited. These thresholds were determined in imaging sessions without light delivery (100 trials) by first calculating the auROC values for each neuron and then pooling the data across all mice. Using a precision-recall curve analysis, we identified the optimal auROC thresholds that best discriminated between excited and other neurons, which were initially identified through statistical testing. We opted for a threshold-based classification approach rather than statistical testing because of the limited number of trials (20 light-OFF or –ON trials) in the session with light delivery, which could have resulted in reduced statistical power.

Neuronal responses to CS were quantified by subtracting pre-CS activity from activity following CS. Statistical comparisons of the responses to CS were performed using a linear mixed-effects model with trial type (high and non) as a fixed effect and random intercepts for mouse, neuron, and trial identities. For the analysis of the imaging session with light delivery, statistical comparisons of responses to CS were performed using a linear mixed-effects model with group (mCherry and eNpHR) and light (OFF and ON) as fixed effects and random intercepts for mouse, neuron, and trial identity. Linear mixed models were fitted using the lmer function in R, followed by Tukey-adjusted post hoc comparisons.

In calcium imaging across 100 trials without light delivery, a linear regression was performed between neuronal responses to CS during all trials and those during high– or non-expression trials. In the imaging experiment with light delivery, linear regression was conducted between the neuronal responses to the CS during the light-OFF and ON trials. Neuronal responses were averaged across trials for each neuron before performing regression. Pearson’s correlation coefficient (r) and slope values were calculated using a linear regression function from the SciPy library.

As a control analysis for Figures S7H–S7J, neuronal activity was compared between trials with high licking frequency and trials without licking during the pre-CS period. For each cell, the increase in activity during the pre-CS period was calculated by subtracting the activity from 6 to 3 seconds before the CS presentation from the activity during the 3 seconds before the CS presentation. Based on the licking frequency during the pre-CS period, trials with the top 10% of licking frequencies were defined as “trials with more licks,” and the first 10 trials with no licking were defined as “trials with no licks.” A linear regression was then performed to examine the relationship between the increase in neuronal activity during the pre-CS period across all trials and those during “trials with more licks” or “trials with no licks.” To analyze the effects of optogenetic inhibition on BLA spontaneous activity, neuronal activity during the 3 s after light delivery was assessed. Statistical comparisons were performed using a linear mixed-effects model, with group (mCherry and eNpHR) and light (OFF and ON) as fixed effects and random intercepts for mouse, neuron, and trial identity.

To analyze the effects of optogenetic inhibition on BLA neuronal population activity, the discriminability of BLA activity between light-OFF and –ON conditions was evaluated using machine learning. Trial-averaged signals from individual neurons were used to train a support vector machine (SVM) classifier with a radial basis function (RBF) kernel. The neurons that were excited by CS in either light-OFF or –ON trials were included in the analysis. The classifier was trained using condition labels (light-OFF or –ON) as target variables. Performance was assessed using repeated stratified k-fold cross-validation (5 folds with 5 repeats) to ensure balanced class distributions across folds. Classification accuracy was computed as the mean accuracy across all cross-validation iterations.

### Statistics and reproducibility

Statistical methods were not applied to predetermine the sample sizes; however, our sample sizes were consistent with those reported in previous studies^56,57^. Data collection and analyses were not blinded to the experimental conditions. Batch and automatic analyses were applied to both the control and experimental groups to minimize the potential for subjective bias. Statistical analyses were performed using GraphPad Prism 10.4.0 (GraphPad Software, La Jolla, California, USA) unless otherwise stated. All raw data processing was performed in Python unless otherwise stated. ANOVA and paired t-tests were used after the distribution of data was visually inspected using histograms to assess normality. Two-sided statistical tests were used, and P < 0.05 was considered statistically significant. A complete list of exact p-values and terms for supporting statistical information is provided in Supplementary Table 1. Data are shown as mean ± SEM, unless otherwise stated. Histological experiments were repeated independently in different mice, with similar results as shown in Figure S1B (n = 5).

### Data availability

The datasets used and analyzed during the current study are available from the corresponding author upon reasonable request.

### Code availability

Custom codes used to analyze data from this study are available upon reasonable request from the corresponding authors.

## ACKNOWLEDGMENTS

We thank Nomura laboratory members for the helpful discussion. We thank K.Shiotani, N.Nomura, and T.Hayashi for technical assistance. We are grateful to Drs. Hiroaki Norimoto, Sho Yamaguchi, and Yasutaka Mukai for their helpful advice on EEG analysis, and to Dr. Hiroyuki Hioki for guidance on the FT-GO immunostaining method. We thank Dr. Karl Deisseroth for providing access to the GCaMP6m, GCaMP6s, ChR2, and eNpHR viral vectors and Vector Core at the University of North Carolina at Chapel Hill and Addgene for distributing these vectors. We also appreciate the assistance from the Research Equipment Sharing Center at Nagoya City University. This work was supported by JSPS KAKENHI Grants (JP23H02787, JP22H05080, JP22K19482, JP21H00296 to HN; JP23K14683 to YM), the JST-FOREST Program (JPMJFR204A to HN), AMED-CREST (JP24gm1510008 to HN), the Suzuken Memorial Foundation (to HN), the Takeda Science Foundation (to HN), Grant-in-Aid for Promotion on Co-Creative Urban Development in Nagoya City University (2401102 to HN), Grant-in-Aid for Outstanding Research Group Support Program in Nagoya City University (24011101), and JST SPRING (JPMJSP2130 to YT). This work was the result of using research equipment shared in MEXT Project for promoting public utilization of advanced research infrastructure (Program for supporting construction of core facilities) Grant Number JPMXS0441500025. We would like to thank Editage (www.editage.jp) for English language editing.

## AUTHOR CONTRIBUTIONS

Y.M., Y.T., K.N., and H.N. designed the study. Y.M., Y.T., K.N., Y.Y., Y.I., R.I., M.O., R.M., and H.N. conducted the experiments. Y.M., Y.T., and H.N. analyzed the data. N.H-I. provided expertise in fiber photometry. Y.M., Y.T., M.M., and H.N. interpreted the data. Y.M., Y.T., and H.N. wrote the manuscript. All authors reviewed and approved the final version of the manuscript.

## ADDITIONAL INFORMATION

Correspondence and requests for materials should be addressed to Hiroshi Nomura.

## FIGURES AND LEGENDS

**Figure S1.**
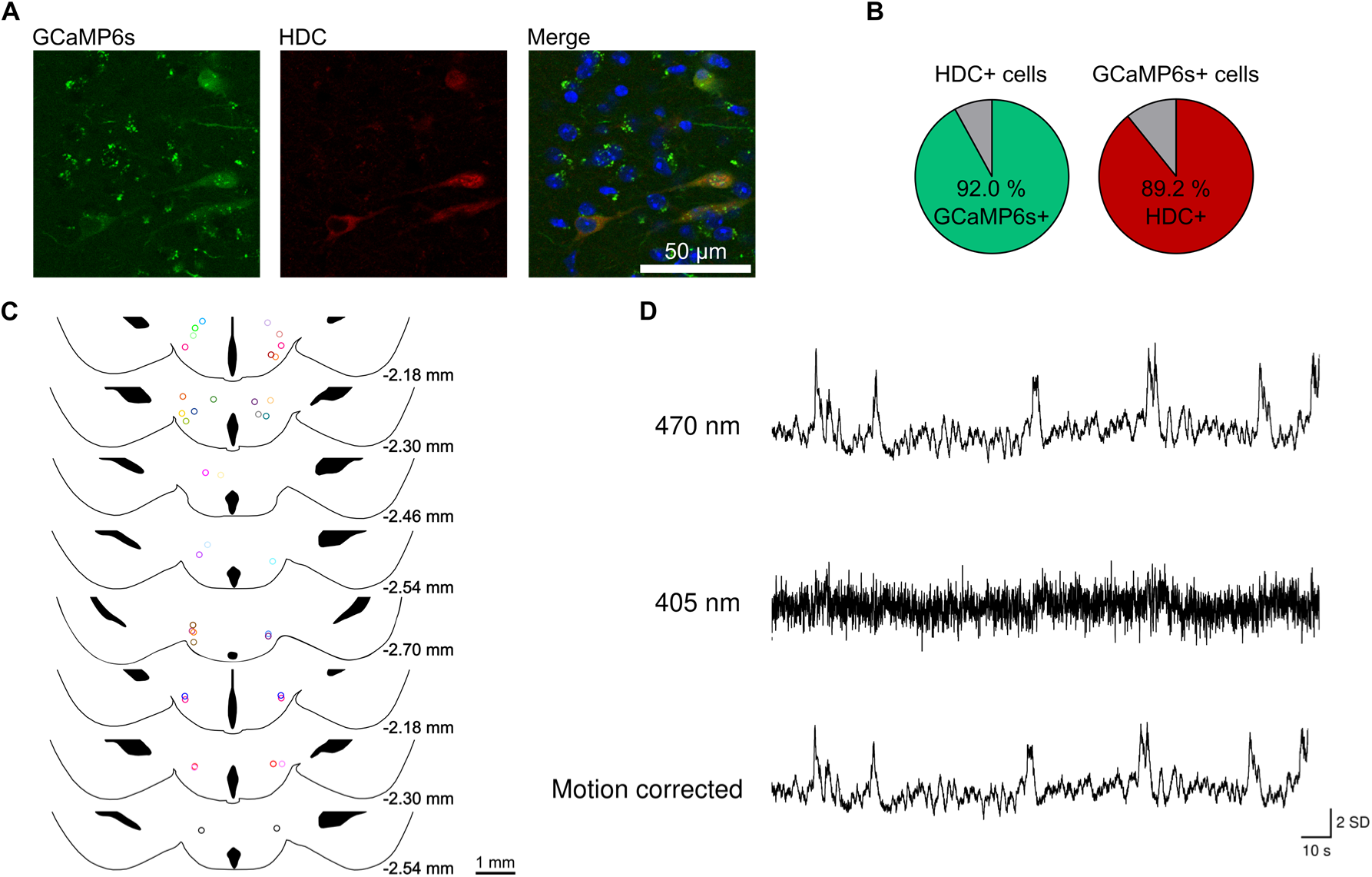
GCaMP6s expression in histaminergic neurons and their calcium activity. (A) Confocal images showing GCaMP6s expression alone, anti-HDC immunostaining alone, and a composite image including DAPI staining in the TMN. (B) Pie charts showing the proportion of cells positive for both HDC and GCaMP6s relative to the total number of HDC-positive cells and GCaMP6s-positive cells. GCaMP6s-positive cells: n = 244 cells, HDC positive cells: n = 237 cells from 5 mice. (C) Locations of the optical fiber tips above the TMN for mice used in Figures 1, 2, and S1–4. (D) Calcium-dependent fluorescence traces (470 nm, top), calcium-independent isosbestic fluorescence traces (405 nm, middle), and motion-corrected traces (bottom).

**Figure S2.**
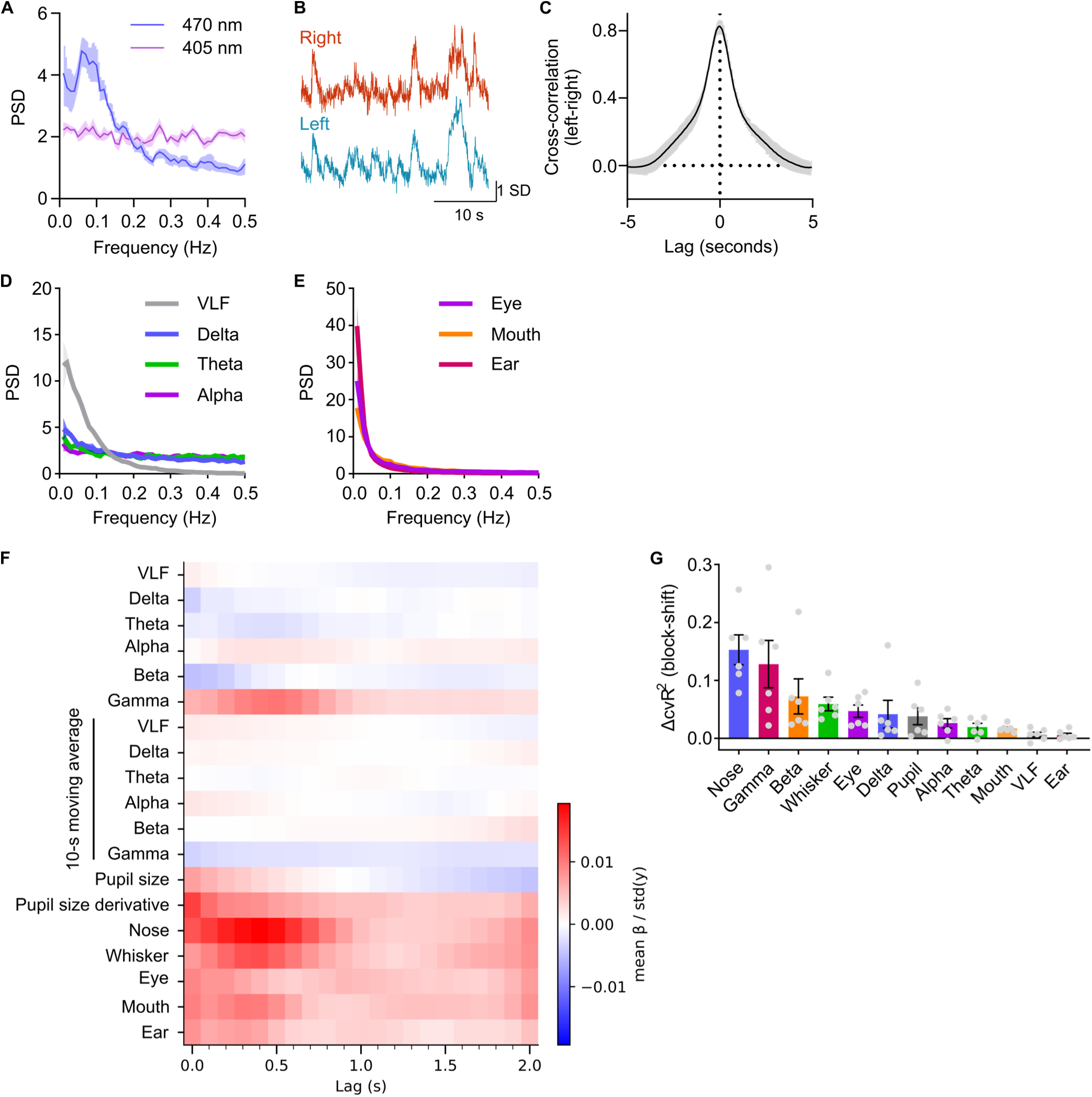
Detailed analysis of spontaneous histaminergic neuronal activity, EEG and facial dynamics. (A) Power spectral density (PSD) of calcium-dependent (470 nm) and calcium-independent (405 nm) fluorescence signals (n = 6 mice). (B) Representative traces showing spontaneous activity in histaminergic neurons of the bilateral TMN. (C) Cross-correlation function between signals from the left and right TMN (n = 5 mice). (D) PSD of each EEG band. (E) For each facial part, power spectra of the top three principal components were summed and then normalized. (F) Kernels of the encoding model to predict histaminergic neuronal activity. Facial movements are represented by the L2-norm of the top three principal components. (G) Decrease in model performance (ΔcvR^2^) after circularly shifting the time series of the indicated feature group. Data are mean ± SEM.

**Figure S3.**
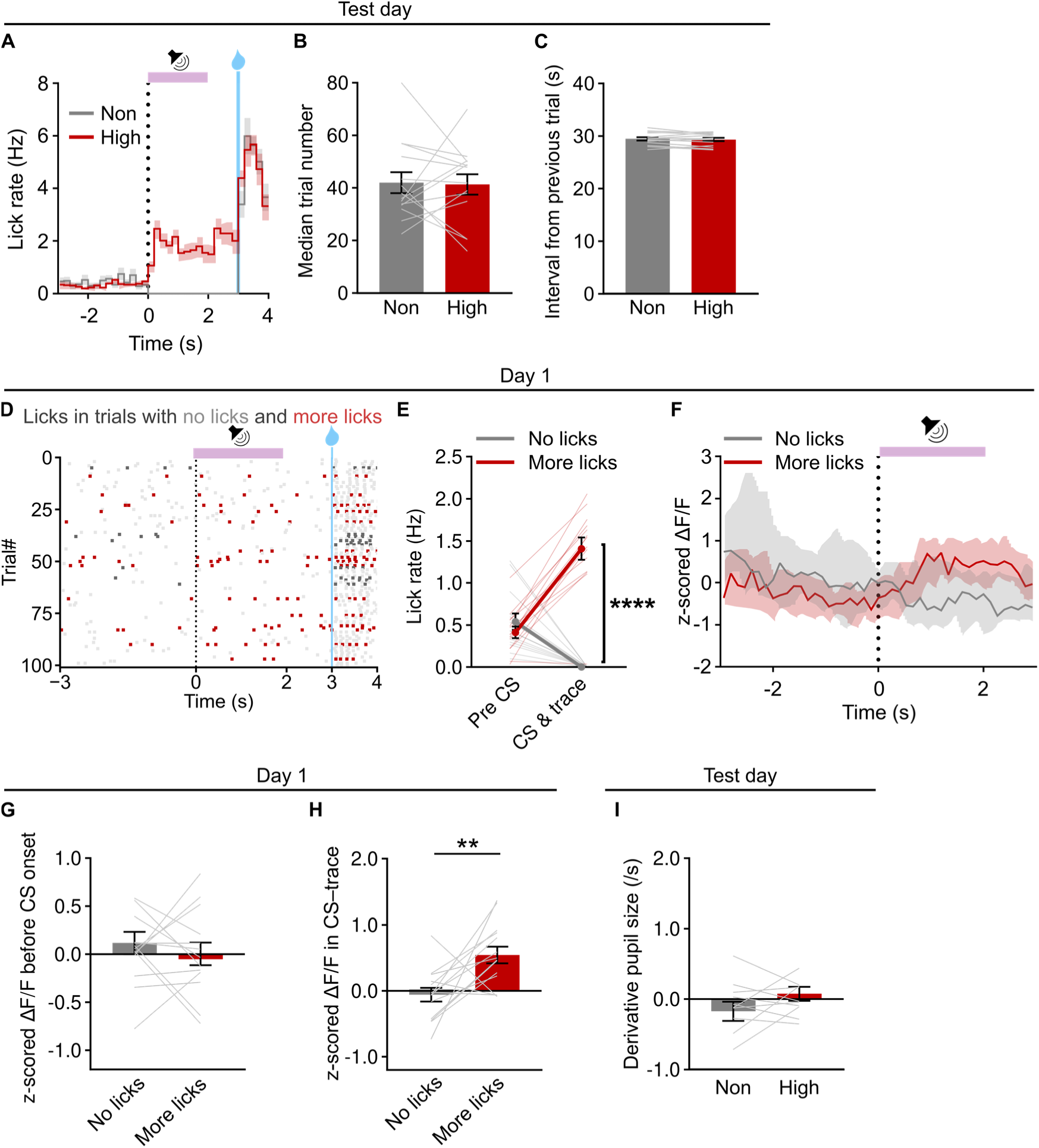
Detailed analysis of data from fiber photometry experiments on the test day and the first day of conditioning. (A–C) Characteristics of high– and non-expression trials on the test day (n = 14 mice). Licking rate around the CS onset in high– and non-expression trials (A). Quantification of the median number of trials (B) and the mean interval between trials (C) in high– and non-expression trials. (D–H) Analysis of behavior and histaminergic neuronal activity in trials with more licks and no licks on the first day of conditioning. (D) Raster plot of licking events from a representative mouse, with color indicating trial types (red: trials with more licks, gray: trials with no licks, light gray: other trials). (E) Licking rates during pre-CS and CS–trace periods in trials with more licks or no licks (****P = 4.24 × 10^-8^, Sidak’s test after repeated-measures ANOVA). (F) The activity of histaminergic neurons around the CS onset in trials with more licks and no licks in a representative mouse. Data presented as median ± top and bottom 25%. (G) The activity of histaminergic neurons before the CS onset (n = 14 mice). Data represented as mean ± SEM. (H) The mean activity of histaminergic neurons during the CS–trace period. (**P = 0.0081, paired t-test). (I) The time derivative of pupil size before CS onset (n = 11 mice). Data are mean ± SEM unless otherwise noted.

**Figure S4.**
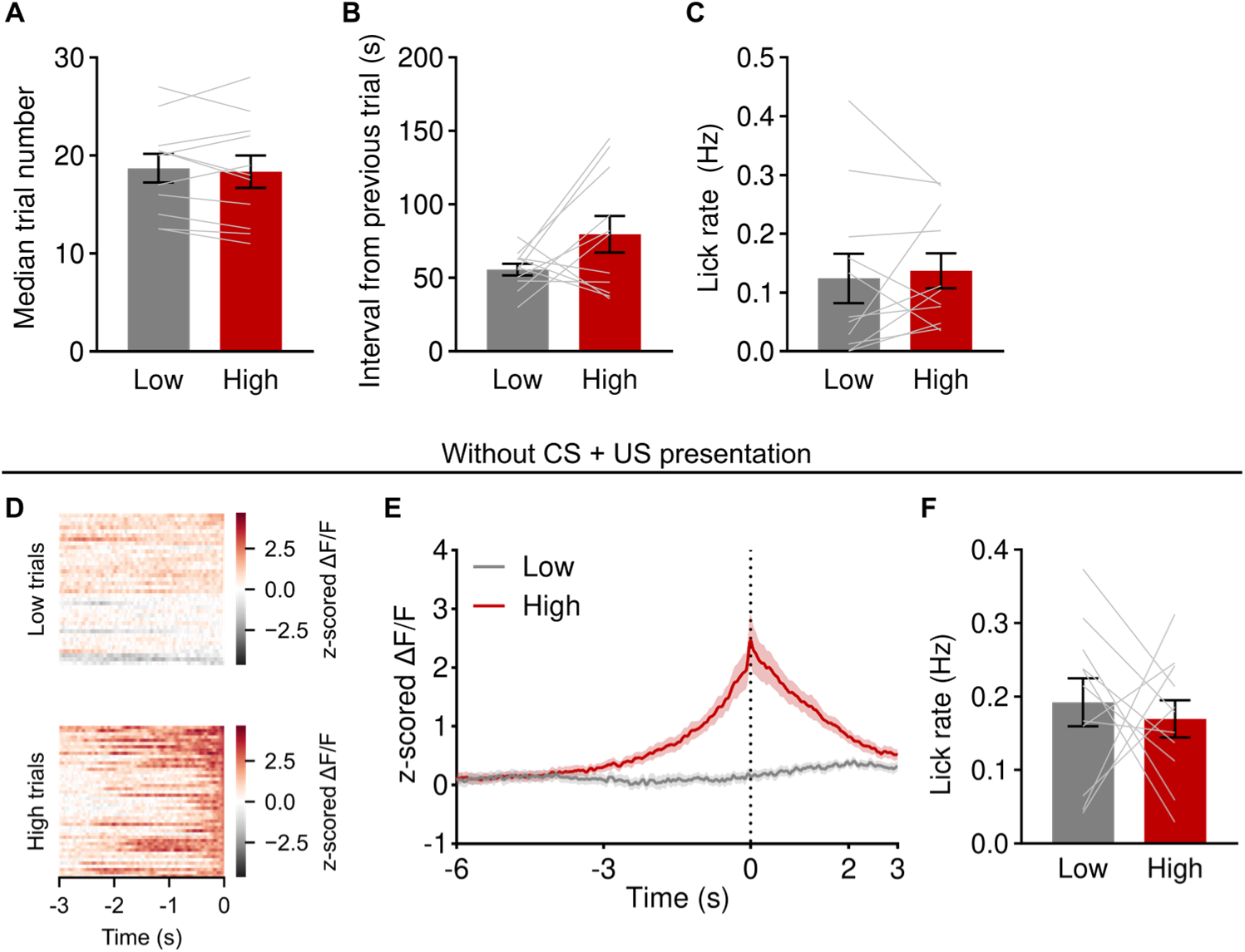
Detailed analysis of closed-loop experiments. (A and B) The median number of trials (A) and mean interval between trials (B) in high and low trials (n = 10 mice). (C) Licking rates measured during the pre-CS period (6 to 3 s before CS onset). (D–F) Analysis of periods with high or low histaminergic neuronal activity in the absence of CS presentation. Heatmap of histaminergic neuronal activity in a representative mouse before detecting high or low conditions (D). Mean activity of histaminergic neurons around the detection of high and low conditions (E) (n = 10 mice). Licking rate during 3 s after detecting high and low activity (F). Data are mean ± SEM.

**Figure S5.**
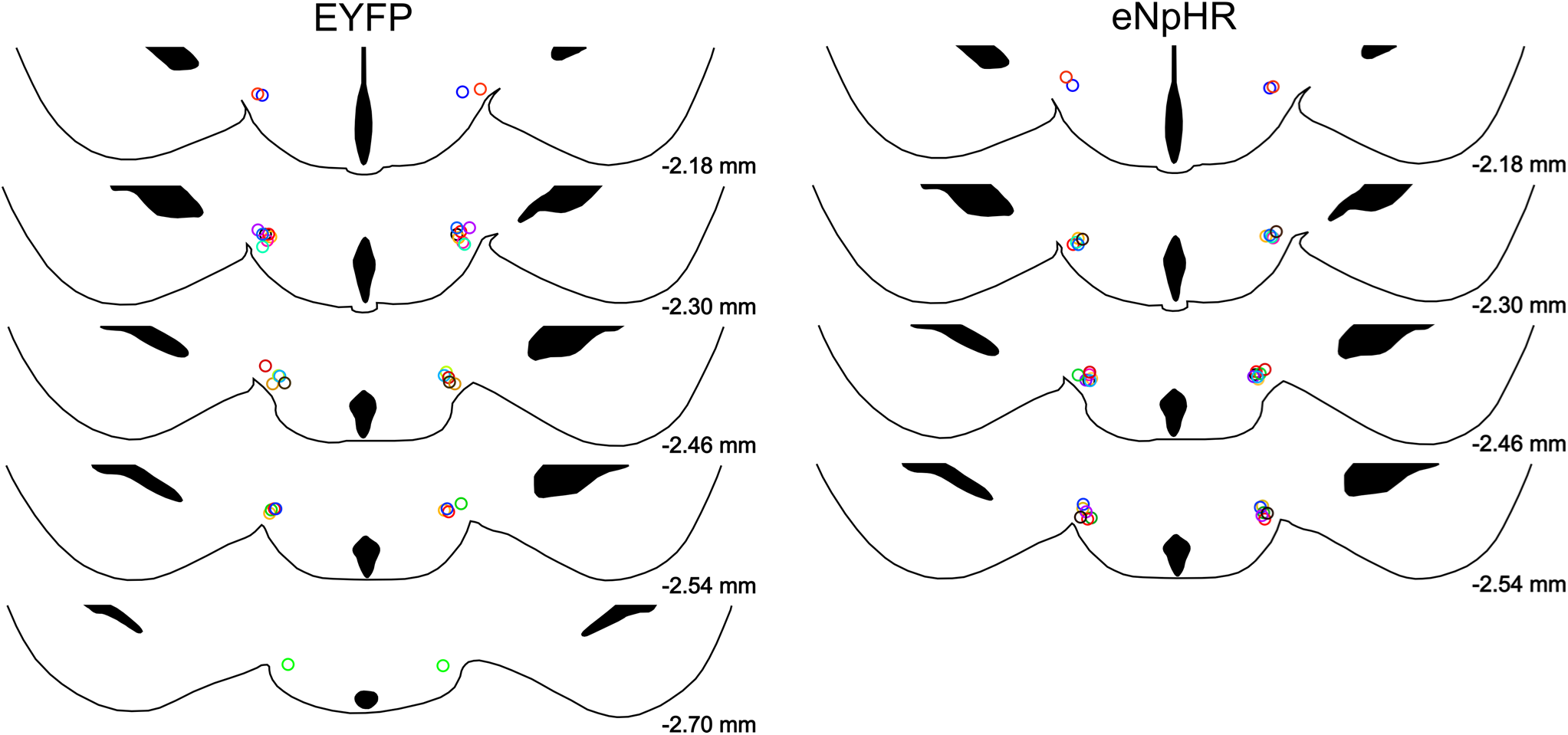
Supplementary data in optogenetic inhibition experiments. Locations of optical fiber tips above the TMN for mice used in Figure 4.

**Figure S6.**
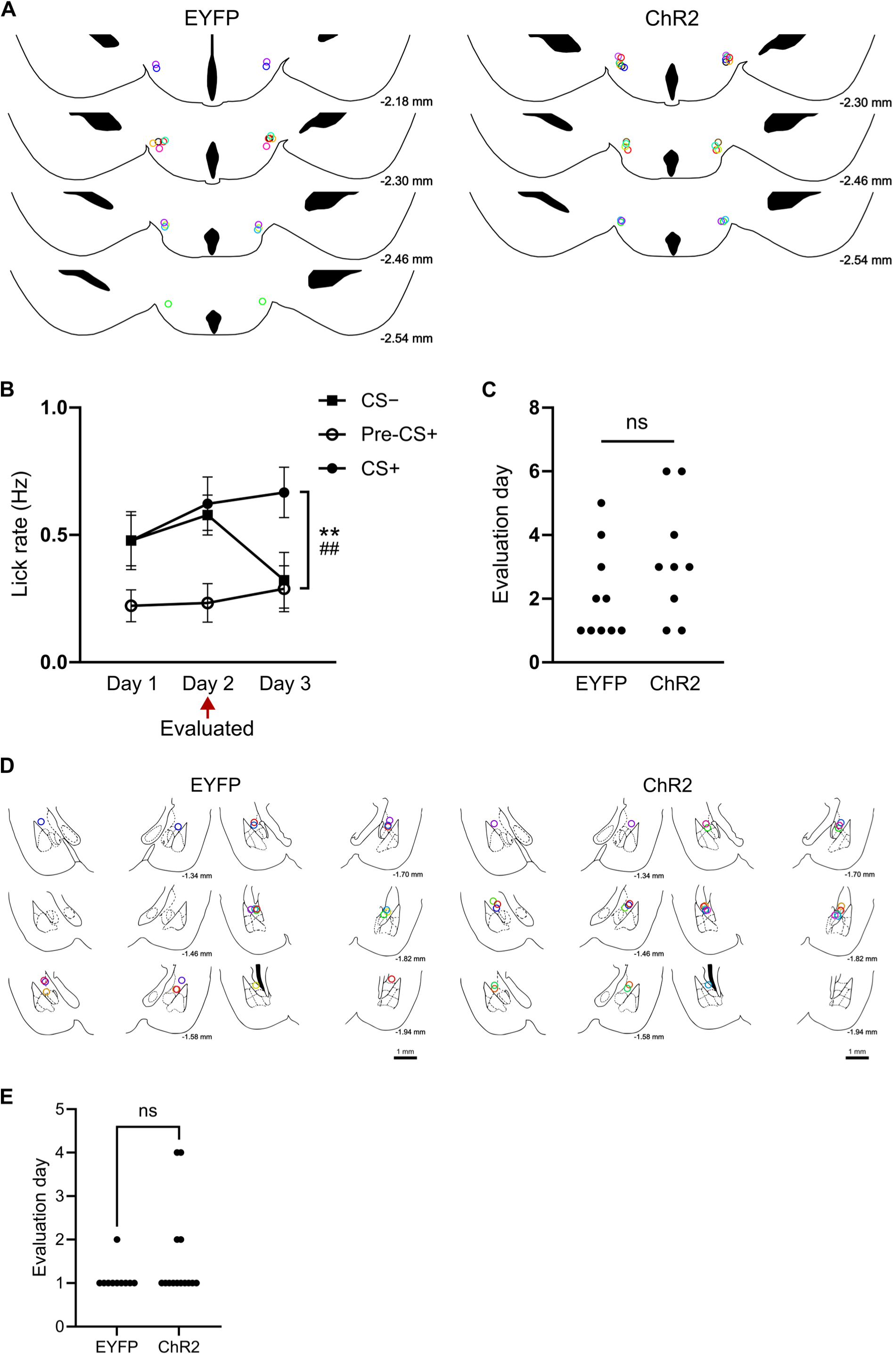
Supplementary data in optogenetic activation experiments. (A) Locations of optical fiber tips above the TMN in mice used in Figure 5. (B) Lick rates during pre-CS+, CS+, and CS− periods on light-OFF trials in a representative EYFP mouse. This example illustrates the approach used to determine the day for evaluating the effect of optogenetic manipulation. This evaluation was performed on the day prior to when lick rates during the CS+ period significantly exceeded those in both pre-CS+ and CS− periods (n = 30 trials; CS+ vs. CS−, **P = 0.00132, Mann–Whitney test; pre-CS+ vs. CS+, ##P = 0.00612, Wilcoxon signed–rank test). Data are mean ± SEM. (C) The designated day for evaluating the effect of optogenetic manipulation shown in Figure 5. (D) Locations of optical fiber tips above the BLA in mice used in Figure 6. (E) The designated day for evaluating the effect of optogenetic manipulation shown in Figure 6.

**Figure S7.**
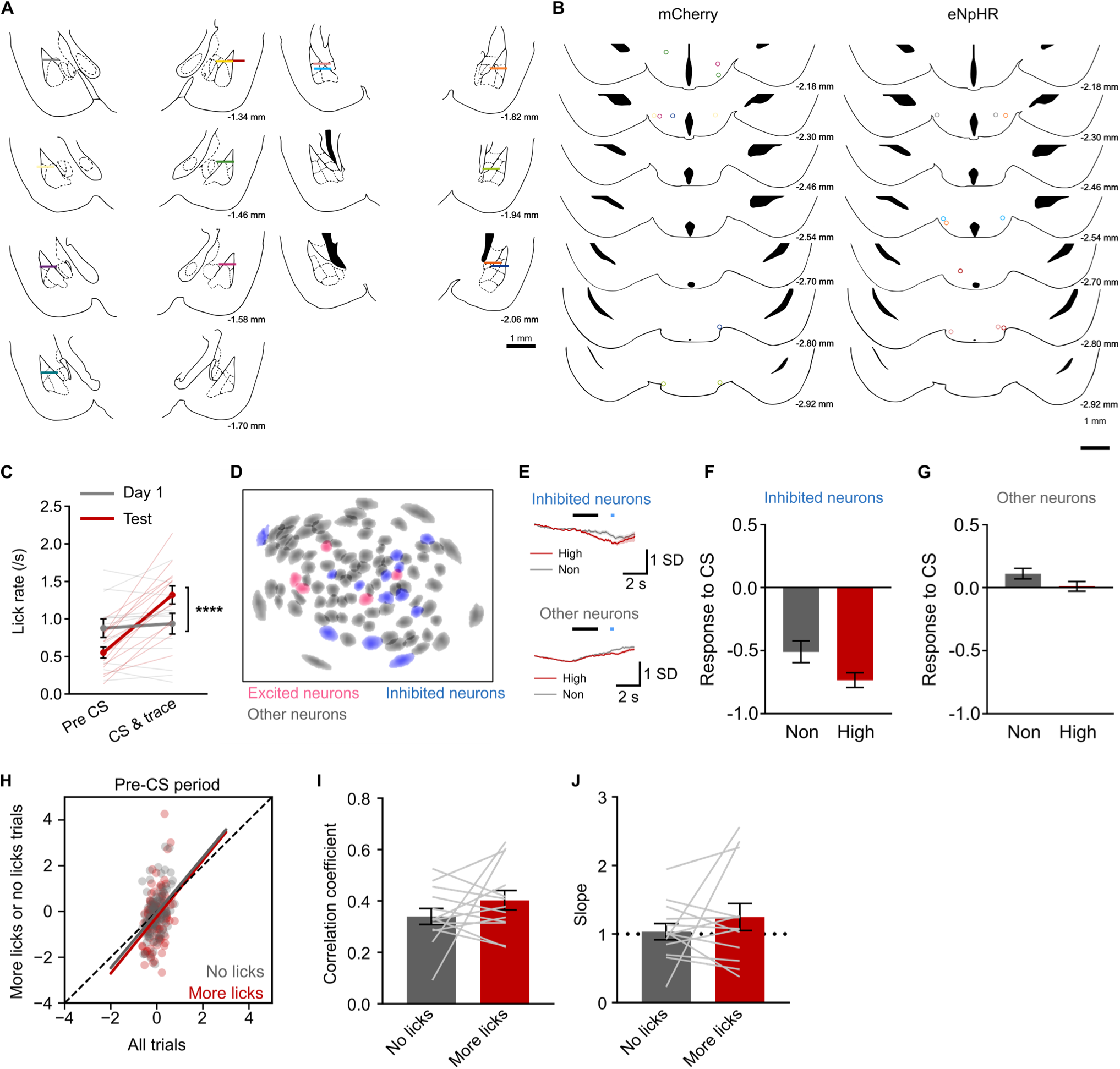
Supplementary data from calcium imaging sessions without optogenetic inhibition. (A) Locations of microendoscope tips above the BLA in the mice used in Figure 7. (B) Locations of optical fiber tips above the TMN in mice used in Figure 5. (C) Licking rates for 3 s during pre-CS period and CS–trace period (Day 1 vs. Test, ****P = 8.56 × 10^-5^, Sidak’s test after repeated measures ANOVA, n = 13 mice). (D) Representative map of neurons in the BLA, with color indicating neuron types. (E) Activity of CS-inhibited and other neurons around CS presentation (n = 145 inhibited neurons and 667 other neurons from 13 mice). (F and G) Neuronal responses of CS-inhibited (F) and other (G) neurons to CS. (H) Scatter plots illustrating neuronal activity during the pre-CS period. The x-axis shows the activity during all trials, and the y-axis represents the activity during trials with more licks or no licks in the pre-CS period. (I and J) Pearson’s correlation coefficients (I) and slopes (J) from linear regression analysis of the scatter plots (n = 13 mice). Data are mean ± SEM.

**Figure S8.**
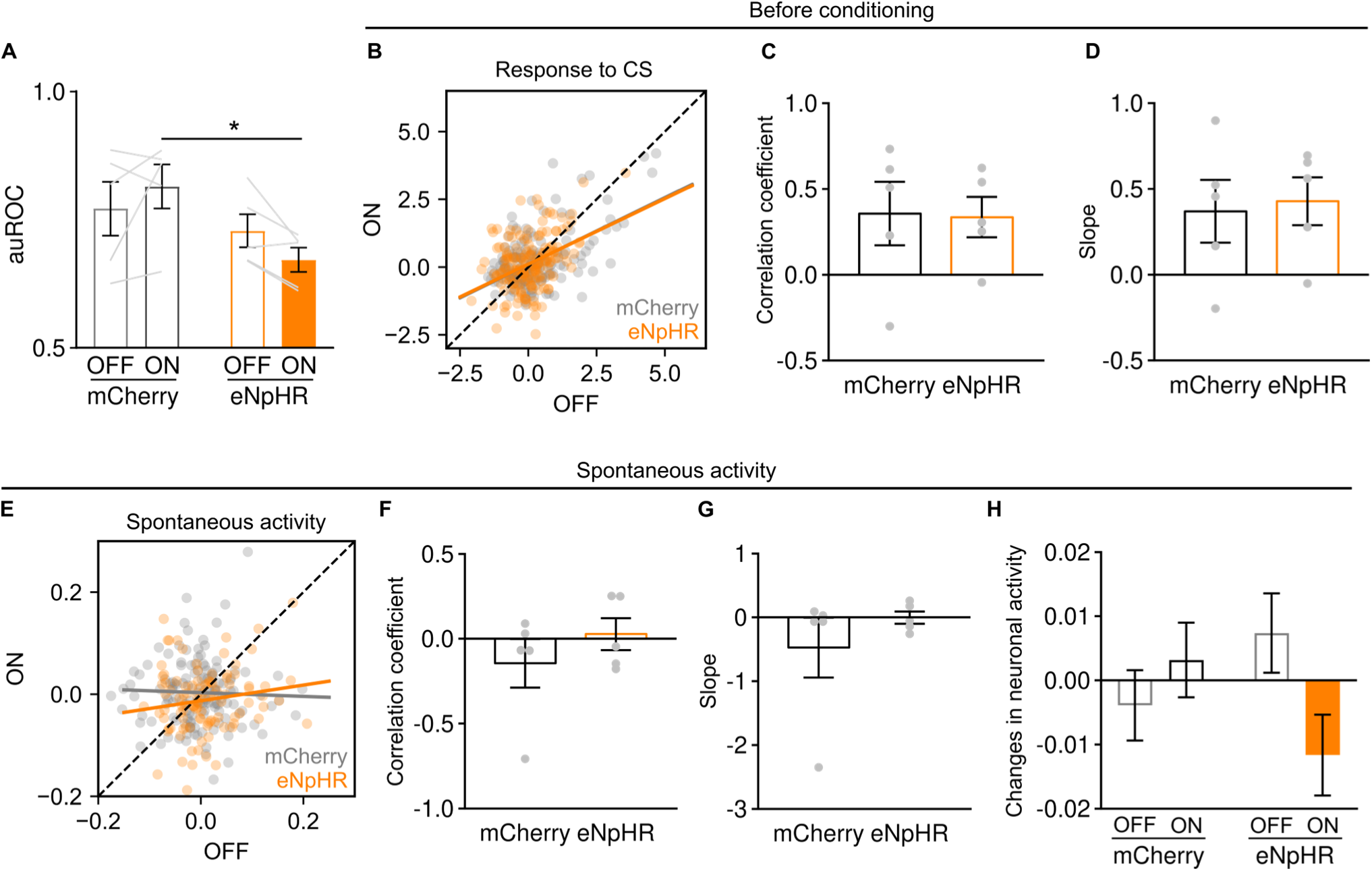
Behavioral and neuronal responses to CS during calcium imaging sessions with optogenetic inhibition. (A) auROC analysis of CS-induced changes in licking rates in mCherry and eNpHR groups. (n = 5 mice (mCherry), n = 5 mice (eNpHR); *P = 0.0407, Sidak’s test; interaction effect between group and laser, F(1, 8) = 4.63, P = 0.0636, two-way repeated-measures ANOVA). (B) Scatter plots illustrating neuronal responses to CS before conditioning. The x-axis shows the response during light-OFF trials, and the y-axis represents the response during light-ON trials, from the mCherry and eNpHR groups. (C and D) Pearson’s correlation coefficients (C) and slopes (D) from linear regression analysis of the scatter plots. (E) Scatter plots illustrating spontaneous neuronal activity for 3 s after 1 s of optogenetic inhibition before conditioning. The x-axis shows the response during light-OFF trials, and the y-axis represents the responses during light-ON trials, from the mCherry and eNpHR groups. (F and G) Pearson’s correlation coefficients (F) and slopes (G) from linear regression analysis of the scatter plots. (H) Changes in BLA spontaneous activity during light-OFF and –ON trials. Data are mean ± SEM.

**Figure S9.**
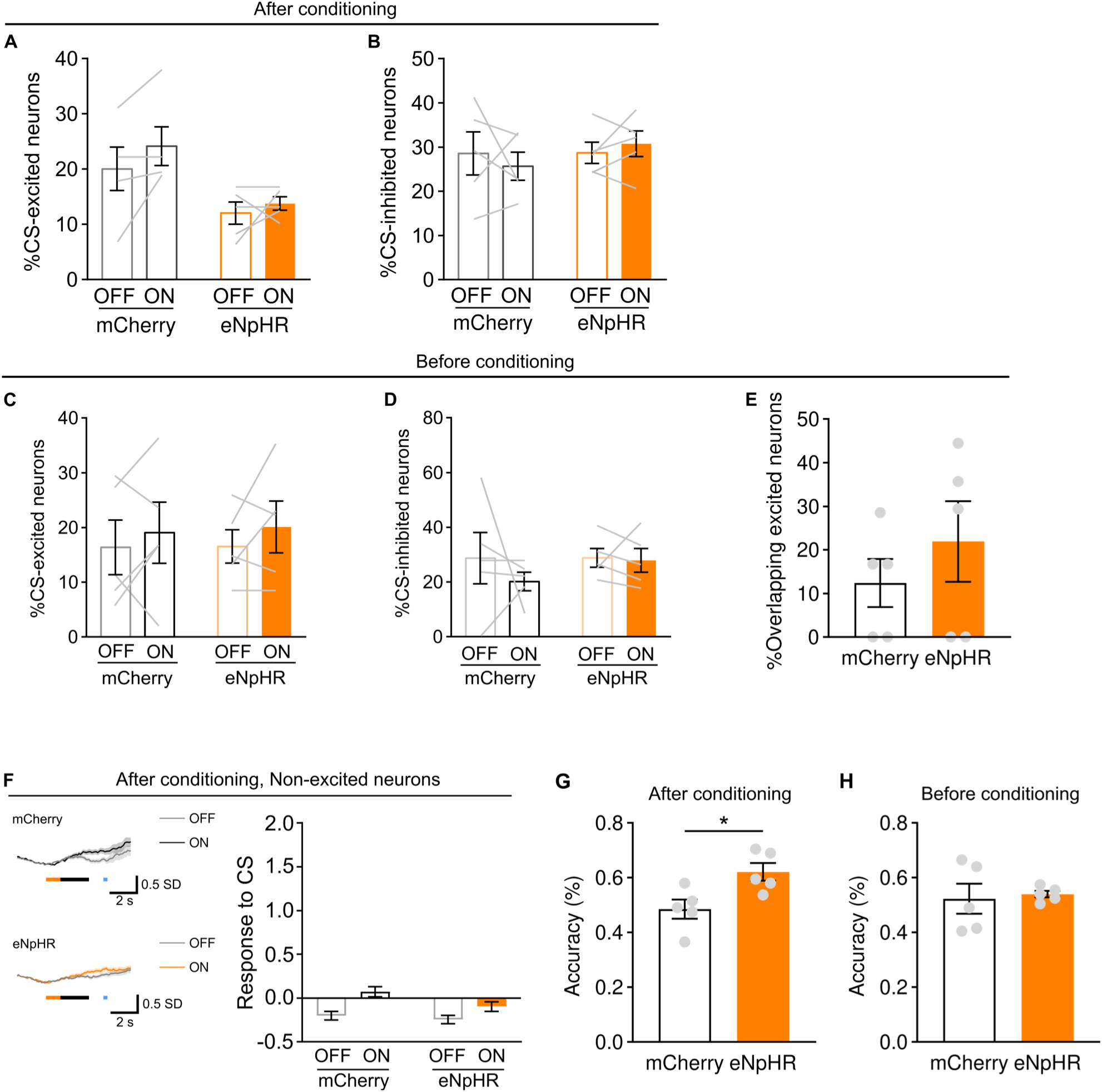
The effects of optogenetic inhibition on neuronal responses to CS before conditioning and spontaneous activity. (A and B) Proportion of neurons excited (A) or inhibited (B) by CS during light-OFF and –ON trials after conditioning. (C and D) Proportion of neurons that were excited (C) or inhibited (D) by CS during light-OFF and –ON trials before conditioning. (E) Proportion of neurons that were excited during both light-OFF and –ON trials among neurons excited in either light condition before conditioning. (F) CS responses of neurons that were not excited by CS during light-OFF trials (mCherry: n = 231 neurons, eNpHR: n = 229 neurons). (G and H) Cross-validated accuracy of two-way decoders discriminating between light-OFF and –ON conditions based on activity data from CS-excited neurons collected after (G) or before (H) conditioning (*P = 0.0215, unpaired t-test). Data are mean ± SEM.

**Figure S10.**
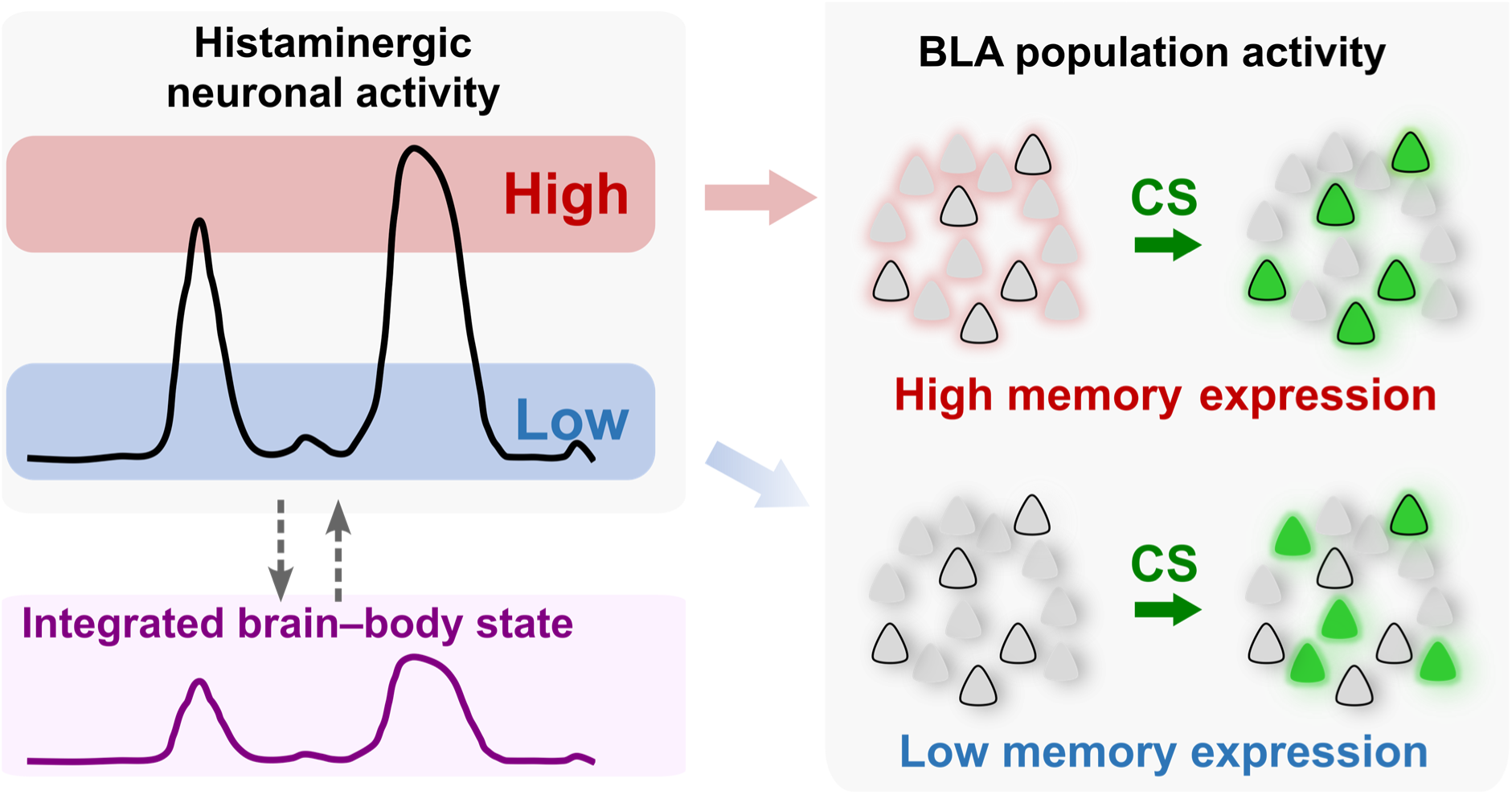
Proposed circuit model for the dynamic modulation of memory accessibility. Ongoing infraslow (0.05–0.1 Hz) activity in TMN histaminergic neurons closely tracks an integrated brain–body state. In our model, elevated pre-CS histaminergic activity primes BLA circuits in a state-setting manner. This priming manifests as a gain-like amplification of the CS-evoked BLA population response with preserved pattern fidelity. In this primed state, the CS more reliably elicits the canonical (operationally defined in this study as the trial-averaged pattern) population response, a pattern associated with successful memory expression. Conversely, when pre-CS histaminergic activity is low, BLA circuits are not primed and the CS elicits a weaker, less faithful population response, yielding diminished memory expression.

